# Neuromuscular, Cardiovascular, and Cognitive Fatigue in Motor Learning: A Systematic Review

**DOI:** 10.1101/2025.09.17.676841

**Authors:** Abdellah Hassar, Jacob Lund, Martin Simoneau, Jason Bouffard

**Affiliations:** Universite Laval; University of Exeter

## Abstract

**Background:** Fatigue is multifactorial and task-dependent, arising from the interplay between performance and perceived fatigability. Fatigue-related changes in sensorimotor control and neural activity may alter skill acquisition and retention.

**Objective:** To synthesize behavioral and neurophysiological evidence on how cognitive, cardiovascular, and neuromuscular fatigue influence motor-skill acquisition, consolidation/retention, and transfer.

**Methods:** A systematic search was conducted within PubMed, Web of Science, and Embase. Thirty-four studies (39 experiments) met inclusion criteria (fatigue effects on motor learning in non-disabled participants). Data were extracted on fatigue protocol, learning taxonomy, learning stage (acquisition vs. retention/transfer), interval, and neurophysiology. Methodological quality was appraised with a tailored Downs & Black checklist.

**Results:** During acquisition, 64% of experiments reported impairments under fatigue, particularly for visuomotor precision and fine force control; 10% found facilitation (often after cardiovascular exertion or high cognitive load); 15% were neutral; 10% inconclusive. Retention was tested in a fresh state and in a fatigue state in 34 experiments and 6 experiments, respectively. Results during retention tests were relatively variable, depending on fatigue protocols. Neurophysiological measures were scarce (EEG/TMS in only four studies), and very few studies considered the effects of personal factors such as age or sex.

**Conclusions:** Fatigue does not uniformly impair motor learning. Neuromuscular fatigue typically hampers acquisition and sometimes retention, while cardiovascular exertion can transiently prime plasticity and enhance consolidation. Evidence for cognitive fatigue is limited and inconsistent. Future work should use longer retention windows, state-matched testing, and integrated neurophysiology to clarify mechanisms and guide training under fatigue.

## 1. Introduction

Fatigue is a task-dependent and multifactorial state that arises from the interaction between performance fatigability, the measurable decline in force or power-generating capacity, and perceived fatigability, the subjective cost of exertion (Enoka & Duchateau, 2016; Russell et al., 2024). These phenomena occur along a continuum shaped by the relative demands placed on cognitive, cardiovascular, and neuromuscular systems. The different dimensions of fatigue interact and influence sensorimotor performance.

Across this continuum, several outcomes are shared by different fatigue states. For instance, cognitive and neuromuscular fatigue can elevate perceived effort, reduce movement precision, accuracy, and impair attentional control (Pageaux & Lepers, 2018). Cognitive fatigue typically emerges with sustained mental activity and is characterized by increased perceived exertion, diminished motivation, and slower executive and attentional processing, processes that are critical for error monitoring and strategy use in motor learning. Neuromuscular fatigue, which develops during repetitive or prolonged muscular activity, reflects peripheral mechanisms, such as metabolite accumulation and impaired excitation–contraction coupling, and central mechanisms, such as reduced voluntary activation and limits in central drive. These mechanisms contribute to reduced force output, movement accuracy and precision, and sensorimotor control, while also increasing perceived effort (Enoka & Duchateau, 2008; Taylor & Gandevia, 2008).

A convergent body of neurophysiological and behavioral evidence indicates that fatigue induced by cognitive, cardiovascular, or neuromuscular exertion can alter sensorimotor control and its neural correlates (Marcora et al., 2009; Pageaux & Lepers, 2018). For instance, prolonged Stroop task performance has been shown to increase intracortical inhibition and reduce motor-evoked potential amplitude, effects that parallel heightened perceived effort and slower corrective responses in visuomotor tracking tasks (Salihu et al., 2022). Similarly, repeated high-intensity muscular contractions lengthen the cortical silent period, reduce voluntary activation, and impair fine motor accuracy and precision during force-matching or pointing tasks (Enoka & Duchateau, 2008; Taylor & Gandevia, 2008). In contrast, a single bout of moderate-to-high-intensity cycling elevates corticospinal excitability, lowers GABA-mediated inhibition, and improves sensorimotor performance, as reflected in faster reaction times and improved manual dexterity (Davranche et al., 2015; Neva et al., 2019). Brief aerobic exercise has also been shown to facilitate long-term potentiation-like plasticity and to transiently enhance the retention of recently acquired motor skills (Neva et al., 2019). A recent meta-analysis of twenty-three studies involving four hundred seventy-four participants confirms that acute aerobic exercise reliably increases transcranial magnetic stimulation indices of corticospinal excitability (Youssef et al., 2024). Taken together, these findings indicate that fatigue and exertion can shift the functional state of the corticospinal system in ways that modify sensorimotor control. Given that the ability to regulate and adapt sensorimotor control is a prerequisite for effective motor-skill acquisition, consolidation, and retention, such physiological and behavioral changes could have downstream consequences for motor learning.

Research on motor learning under fatigue conditions has produced relatively heterogeneous findings (Taylor, 2018). For example, moderate neuromuscular fatigue has been shown to alter the internal model acquired during a force field adaptation task (Takahashi et al., 2006). On the other hand, exhaustive running led to improved visuomotor adaptation during a trajectory tracking task (Mierau et al., 2009). These contradictory results may arise from differences in the type of fatigue induced, the characteristics of the task, or the mechanisms of motor learning engaged. Motor learning is not a unitary process; it encompasses partly dissociable mechanisms, such as error-based adaptation, use-dependent plasticity, reinforcement learning, and off-line skill consolidation, which act on different timescales and are supported by distinct neural substrates (Krakauer et al., 2019)

The objective of the current systematic review is to determine how cognitive, cardiovascular, and neuromuscular fatigue influence motor-skill acquisition, consolidation, retention, and transfer across different types of motor learning tasks. This review critically examines two complementary streams of evidence: (i) behavioral outcomes that quantify motor performance during learning with and without fatigue, and (ii) neuroimaging and neurophysiological measures that reveal the mechanisms underlying the interactions between fatigue and motor learning.

We hypothesize that neuromuscular fatigue will impair motor-skill acquisition and final performance in tasks requiring high force or power, due to its negative effects on contractile capacity and corticospinal excitability. Cardiovascular fatigue may enhance early skill acquisition, consistent with an aerobic priming effect reported in several studies(Lehmann et al., 2022; Moriarty et al., 2022). Cognitive fatigue is expected to have little effect on initial error-based adaptation but to reduce retention when mental load is high, with this impact related to changes in cortical excitability and attentional control.

## 2. Method

### 2.1. Study Design

This systematic review was carried out following the Preferred Reporting Items for Systematic Reviews and Meta-Analyses (PRISMA) guidelines. The protocol for the review was registered in the International Prospective Register of Systematic Reviews (PROSPERO, Registration Number CRD42023394696).

### 2.2. Search Strategy

A systematic search was conducted in PubMed, Web of Science, and Embase from their inception until November 2022. The search was developed based on standardized vocabulary (e.g., Medical Subject Headings) and keywords for two general categories: fatigue and motor learning. Boolean operators (OR and AND) were employed to combine standardized terms and keywords.

### 2.3. Inclusion and Exclusion Criteria

We included studies involving participants without any disabilities that investigated the effects of fatigue, whether induced by motor or cognitive effort, on motor learning. To be eligible, the assessment of fatigue’s impact on motor learning had to be the primary objective of the study. Consequently, we excluded studies in which fatigue may have affected motor learning but was not the main focus (e.g., studies on the effects of acute exercise on motor learning (Marin Bosch et al., 2020) or massed vs. distributed practice (Kwon et al., 2015). Conference abstracts were excluded. Study selection was conducted using Covidence software in a two-phase process. First, titles and abstracts were independently screened by two reviewers (AH, JL) based on predefined inclusion and exclusion criteria. Then, potentially eligible full-text articles were assessed independently by the same two reviewers. Any disagreements or uncertainties were resolved through discussion or by consulting a third reviewer (JB).

### 2.4. Quality Evaluation

The methodological quality of included studies was assessed using a customized version of the Downs and Black checklist (Downs & Black, 1998), a standardized tool for evaluating the methodological quality of randomized and non-randomized studies. In this review, the checklist was adapted to address the specific methodological aspects relevant to research on fatigue and motor learning. Five additional items were included to evaluate: (1) the description and appropriateness of the fatigue-inducing task, (2) the description and appropriateness of the motor learning task, (3) comparability between fatigued and control groups, (4) baseline comparability prior to training, and (5) clarity in the reporting of results related to fatigue-induced effects on motor learning. In the resulting checklist, each of the 28 items evaluates a specific methodological criterion (e.g., clarity of hypothesis, validity of outcomes, adequacy of statistics). The total score (0–28) is the sum of these items: a higher score indicates stronger methodological quality, while a lower score highlights methodological weaknesses. Each study was independently rated by two reviewers (AH, JL), with discrepancies resolved through discussion or, if necessary, consultation with a third reviewer (JB).

### 2.5. Data Extraction

Data were extracted systematically into a structured extraction table. Extracted data included study characteristics, participant demographics, fatigue protocol details, motor learning task descriptions, retention interval, outcome measures (grouped into performance and neurophysiological), and results from both acquisition and retention phases with and without fatigue. One author (AH) extracted all data and reviewed it with a second author (JB). Disagreements were resolved by consensus.

### 2.6. Data Synthesis

The studies were first categorized according to the type of fatigue induced: cognitive fatigue (resulting from sustained mental activity), local neuromuscular fatigue (resulting from repetitive or prolonged activation of muscle groups involved in the motor learning task), or cardiovascular fatigue (resulting from prolonged whole-body aerobic or anaerobic exertion). A single experiment could involve more than one type of fatigue; in such cases, all relevant categories were noted (Figure 1).

**Figure 1.**
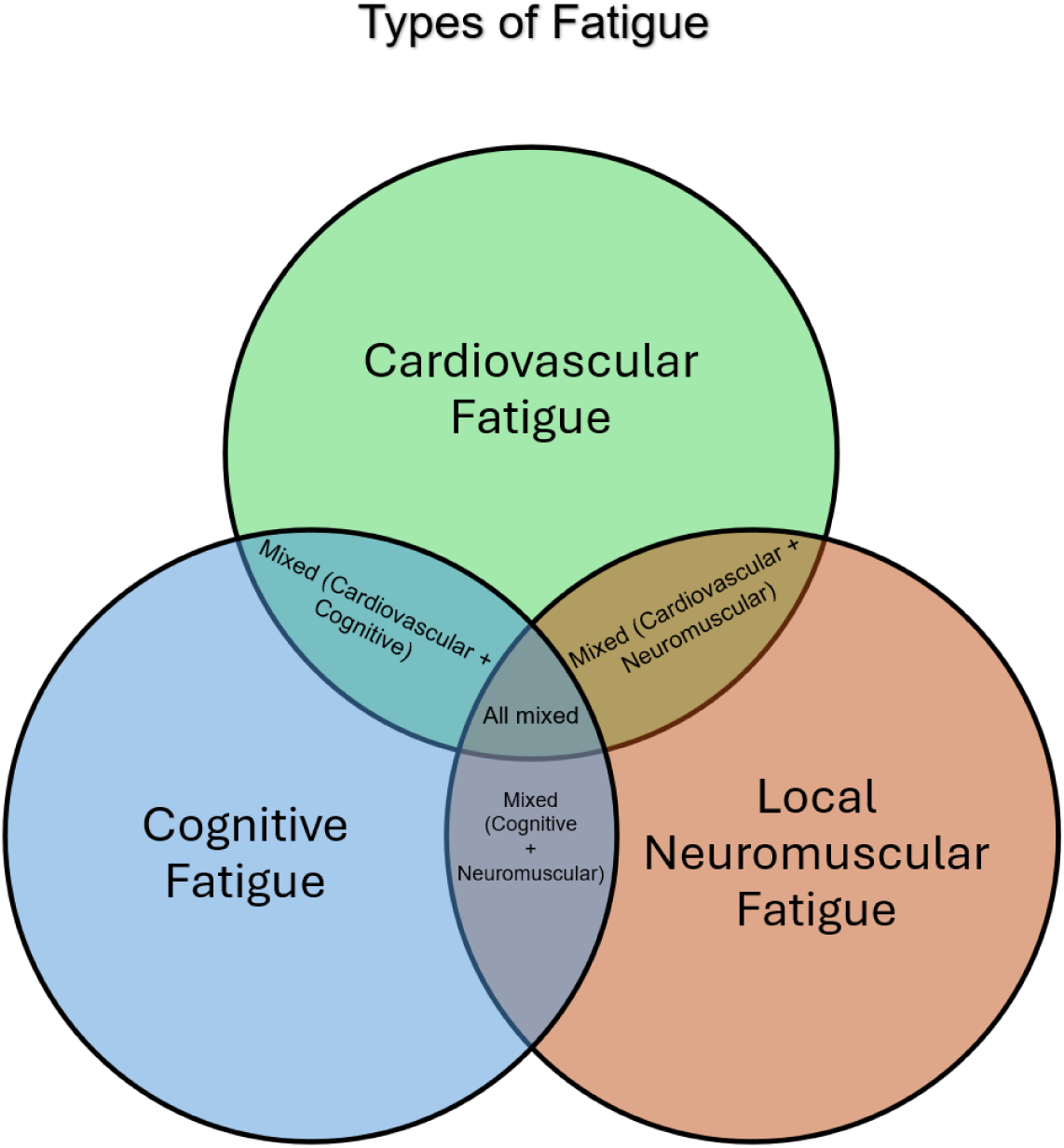
Conceptual Framework of Fatigue Types in Motor Learning.

Subsequently, studies were organized using the motor learning taxonomy of Krakauer et al. (2019), which distinguishes between discrete and continuous *de novo* learning (acquisition of entirely new movement patterns), discrete and continuous sequence learning (learning and optimization of a specific sequence of actions), motor adaptation and learning of motor acuity tasks (Figure 2). As for fatigue protocols, motor learning tasks could be categorized into multiple categories. For each study, results were extracted for the stage of motor learning assessed: acquisition (performance changes during practice) and retention/transfer (the ability to reproduce or generalize the skill after a delay, in the same or a different context, tested with or without fatigue). Retention tests were considered as reflecting both consolidation (long-term memory stabilization) and recall processes.

**Figure 2.**
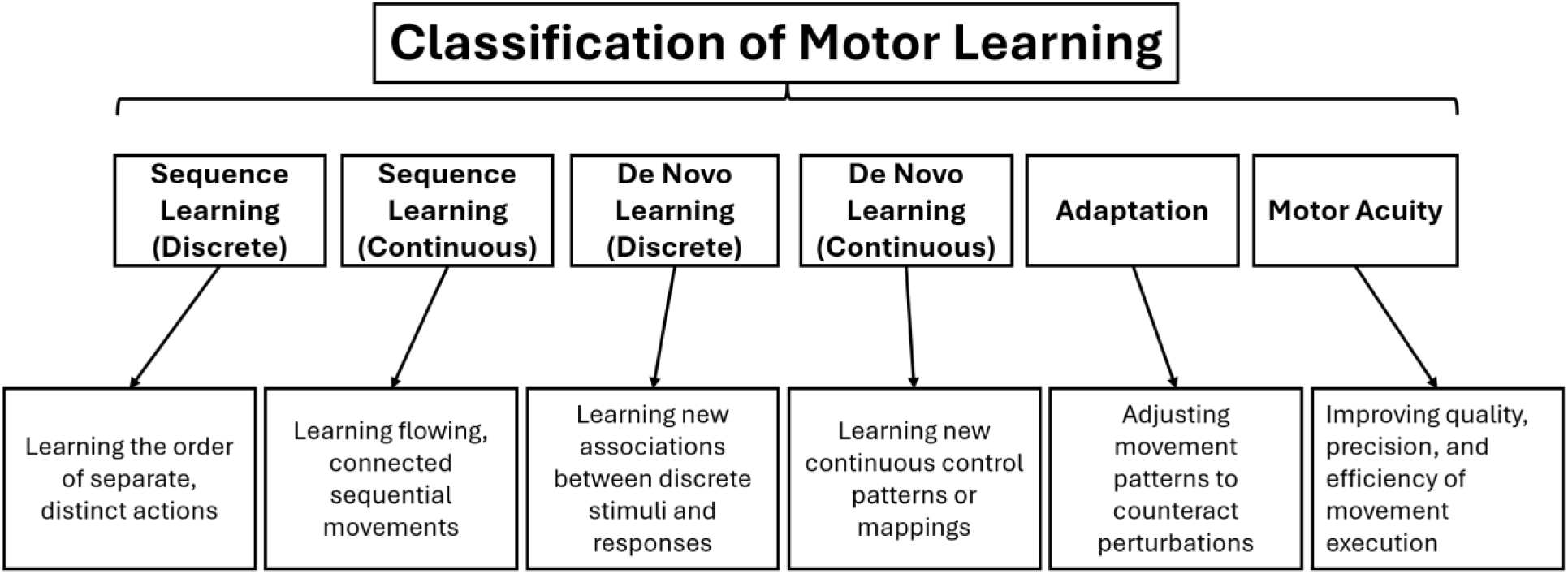
Conceptual classification of motor learning. This framework illustrates six categories of motor learning adapted from Krakauer et al. (2019).

This multilayered classification enabled a more nuanced comparison of findings across protocols and study designs. By integrating fatigue type, retention context, and both the learning type and stage of motor learning, we aimed to identify potential moderators influencing the effects of fatigue on motor learning outcomes and to guide the development of more effective training and rehabilitation strategies.

## 3. Results

The literature search strategy yielded 2,597 titles and abstracts after removing duplicates. Following the application of inclusion and exclusion criteria and the addition of 5 studies through manual citation tracking, 34 studies were included (Figure 3). Results of a total of 39 experiments were reported within these articles.

**Figure 3.**
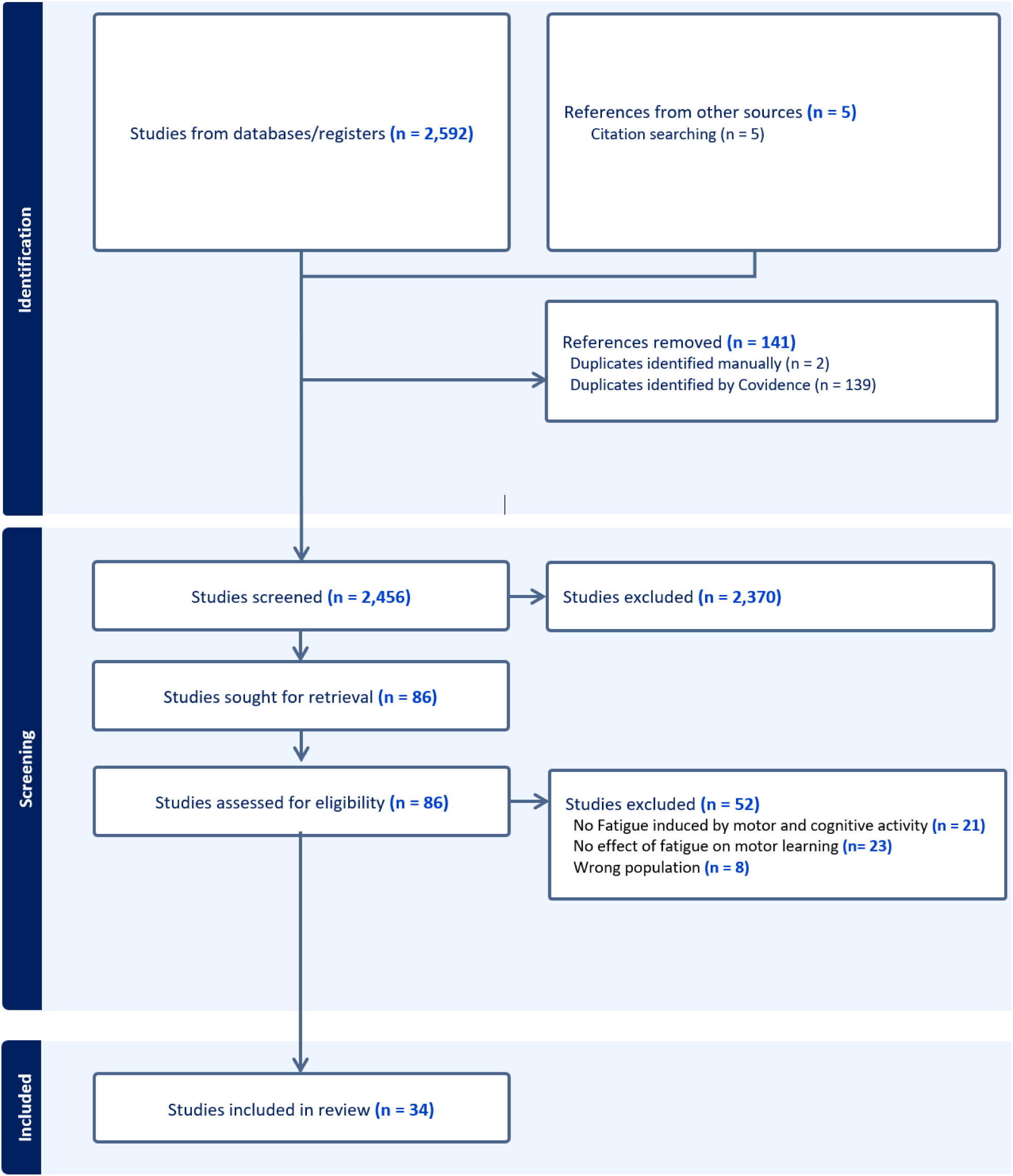
PRISMA flow chart for the identification, screening, and inclusion of articles. n represents the number of texts.

### 3.1. Quality Assessment

The methodological quality of the 34 studies was moderate overall (Table 1). Total Downs and Black scores ranged from 5 (Stockard, 1974) to 18 (Zabihhosseinian et al., 2020; Zabihhosseinian et al., 2021), with a mean score of 12.6/28. Most studies clearly described their hypotheses and outcome measures, applied appropriate statistical analyses, and reported variability measures. However, recurrent weaknesses included insufficient blinding, limited detail on participant selection and follow-up, and uncertainty about outcome validity and reliability. Publication year explained 35% of the variance in quality scores, showing that methodological rigor has improved over time, though quality remained heterogeneous across periods.

**Table 1:**
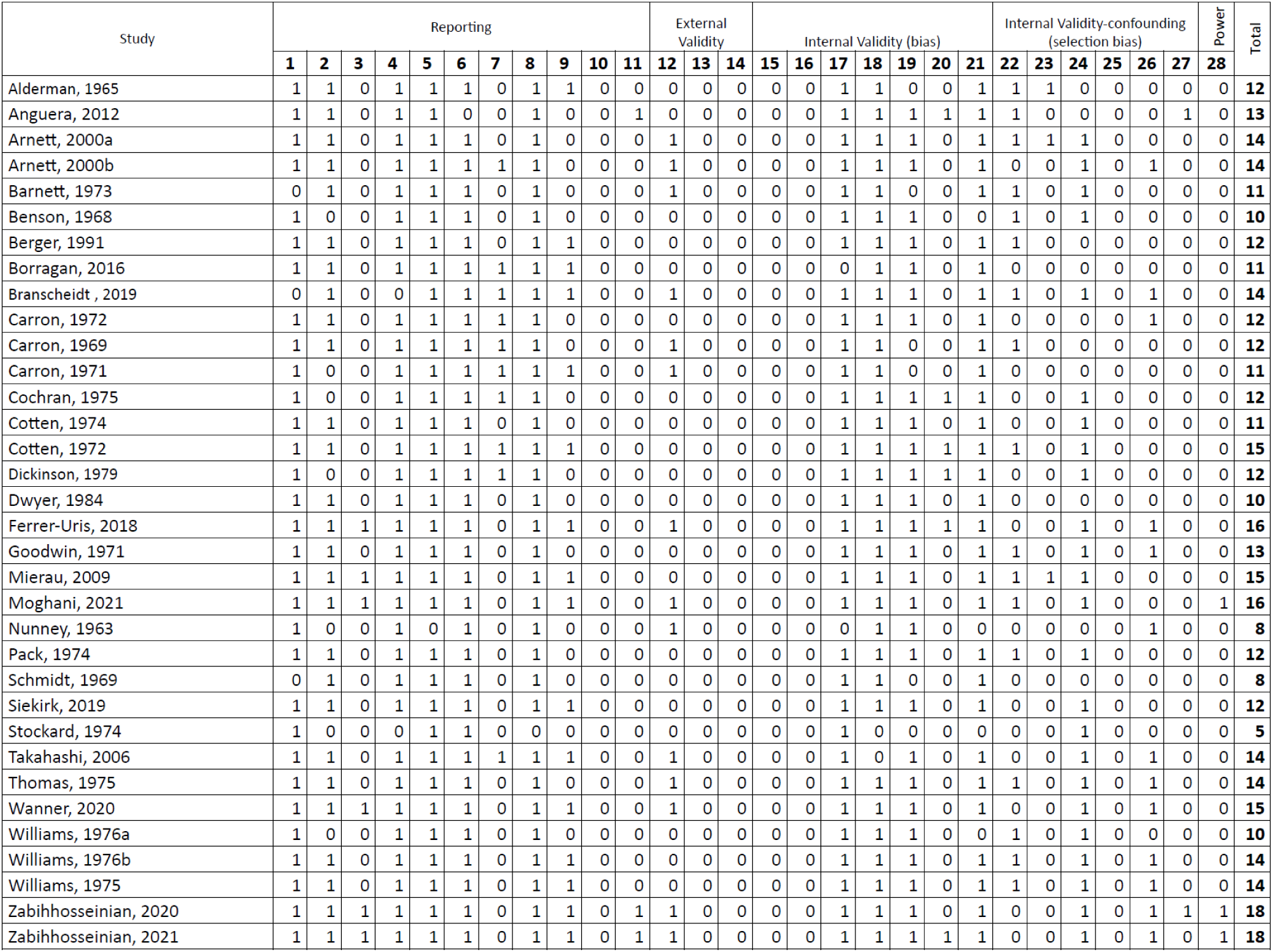
Downs & Black Checklist – Methodological Quality Assessment. Criteria: 1. Hypothesis | 2. Outcomes | 3. Participants | 4. Fatigue task | 5. Motor task | 6. Group comparability | 7. Baseline comparability | 8. Results clarity | 9. Variability reported | 10. Adverse events | 11. Follow-up | 12. p-values | 13. Representativeness (invited) | 14. Representativeness (accepted) | 15. Blinding subjects | 16. Blinding assessors | 17. Data dredging | 18. Follow-up duration | 19. Statistical tests | 20. Compliance | 21. Validity/reliability | 22. Same source population | 23. Recruitment period | 24. Randomisation | 25. Allocation concealment | 26. Confounding adjustment | 27. Attrition | 28. Statistical power.

### 3.2. Characteristics of the studies

The included experiments predominantly recruited relatively homogeneous samples of healthy young adults. When age was reported, mean values typically ranged from 19 to 27 years, although several studies provided only general descriptors (e.g., “college students”) or did not report age at all. At the participant level (Figure 4-A), women represented about 36% of the pooled sample (n = 598 of 1,660), while men accounted for 64% (n = 1,062). At the study level (Figure 4-B), 15% of the experiments tested only women (n = 356 participants), 49% tested only men (n = 805 participants), and 36% included mixed-gender cohorts (n = 499 participants). Notably, only one study (Arnett et al., 2000b) conducted formal sex-specific analyses to directly compare responses between women and men.

**Figure 4.**
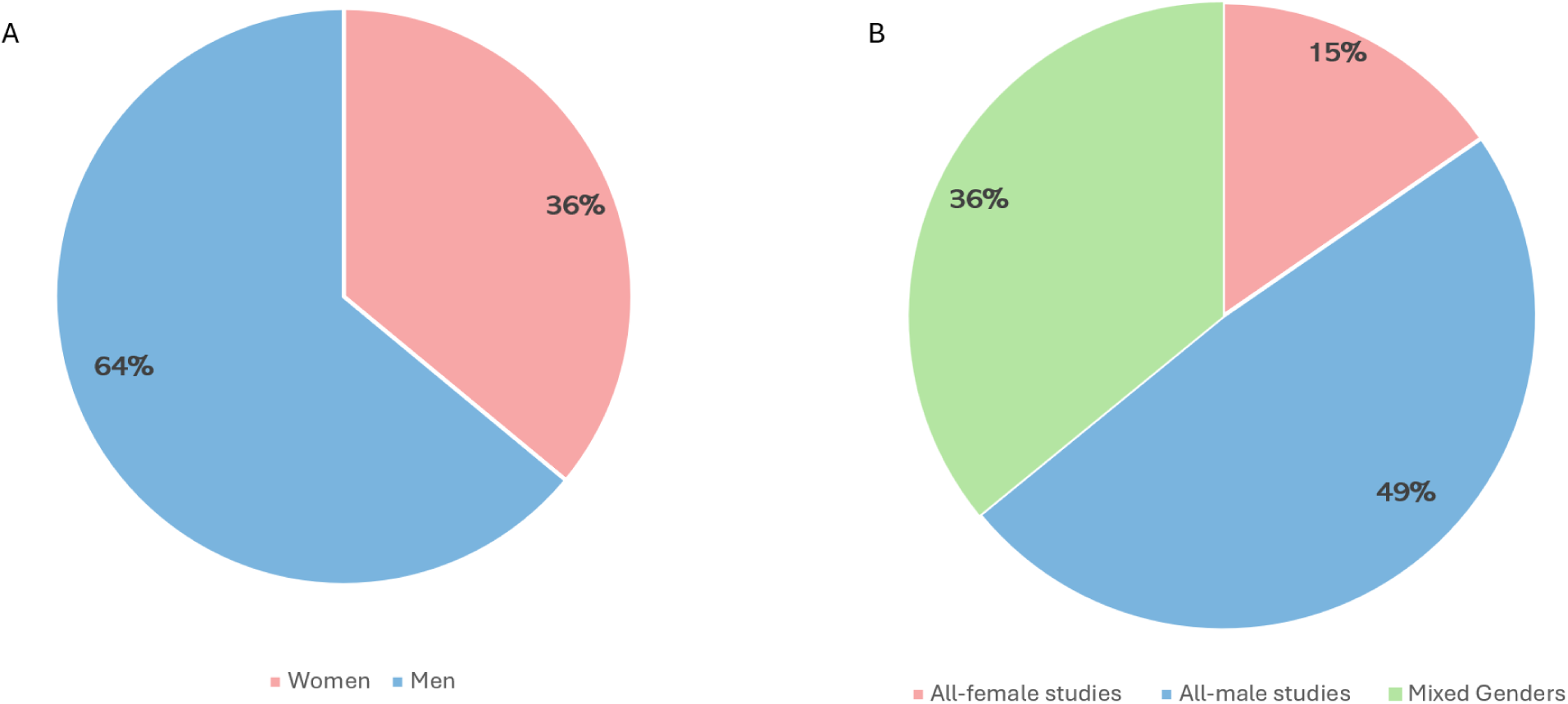
Sex Representation in in included Studies. (Panel A: Distribution of participants; Panel B: Distribution of study cohorts).

Fatigue-induction protocols were highly heterogeneous (Supplementary Table 1). Most experiments induced localized neuromuscular fatigue with a concomitant cardiovascular load (n = 24; e.g., hand-ergometer exercise at ≥ 85 % VO₂ max; Carron (1969) or without any cardiorespiratory component (n = 9; e.g., a sustained pinch at 40 % MVC; (Branscheidt et al., 2019)). Three experiments produced fatigue through cardiovascular exercise performed by a body segment not involved in the motor-learning task, thus avoiding local neuromuscular fatigue, such as a shuttle-run preceding an upper-extremity skill (Mierau et al., 2009). Finally, three experiments elicited fatigue via cognitively demanding paradigms (e.g., Stroop or working-memory tasks; (Anguera et al., 2012; Borragan et al., 2016; Khojasteh Moghani et al., 2021).

Motor learning tasks encompassed the full spectrum described by Krakauer et al. (2019), and many tasks included more than one type of motor learning. Three experiments assessed adaptation paradigms (e.g., cursor rotations and velocity-dependent force fields) that required recalibration of an existing controller. De novo learning tasks that demanded the construction of a novel visuomotor controller were used in 13 experiments, including eight continuous protocols such as pursuit-rotor tracking or sinusoidal tracing and three discrete mappings such as mirror-toss. Ten experiments required participants to acquire the temporal order of discrete actions (discrete sequence-learning; e.g., serial reaction-time task) and 21 involved the production of a sequence of motor actions during a continuous task (continuous sequence-learning; e.g., ladder climbing). Finally, almost all experiments evaluated motor-acuity or skill-refinement protocols (n = 35). All 39 experiments documented performance changes during the acquisition phase, and retention was assessed in 36 protocols, of which two tested retention under fatigue, 30 after complete recovery, and four in both conditions. Retention intervals varied substantially, ranging from immediate retests conducted within ten minutes of practice in eight experiments to multi-day follow-ups extending from 48 hours to two weeks in 15 experiments. Across studies, the median retention interval was 24 hours (Figure 5), highlighting the predominance of short-term assessments. Studies using objective neurophysiological measures were scarce, as only four studies employed techniques such as electroencephalography (EEG; e.g., Mierau et al. (2009)) or transcranial magnetic stimulation (TMS; e.g., Branscheidt et al. (2019), Exp. 3; Zabihhosseinian et al. (2021)), underscoring the limited mechanistic evidence available to complement behavioral outcomes.

**Figure 5.**
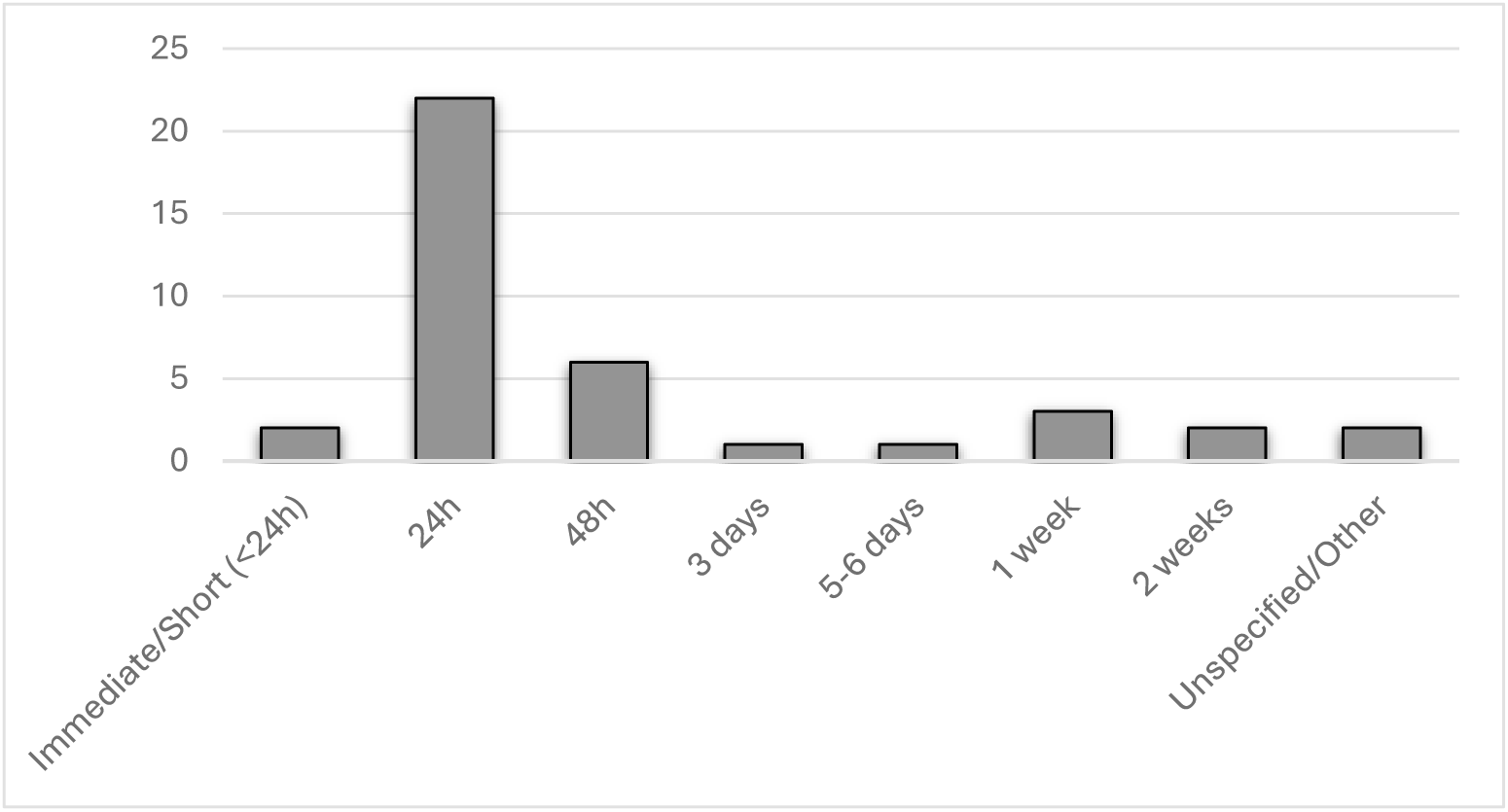
Distribution of retention intervals in fatigue and motor learning studies.

### 3.3. Effects of Fatigue on Motor-Skill Acquisition

On Day 1 of training, 64% of the experiments (25/39) showed a detrimental impact of fatigue on task acquisition, typically expressed as slower learning rates, higher error scores, or a deterioration of the speed–accuracy trade-off (Table 2). Impairment was particularly evident in tasks requiring fine force modulation or precise visuomotor coordination. For instance, in the sequential pinch-force task of Branscheidt et al. (2019), Exps. 1–3), fatigue clearly reduced performance, consistent with tasks categorized as motor-acuity and discrete sequence learning in Krakauer’s taxonomy. By contrast, in their ten-element sequence paradigm (Exp. 4), which required key pressing without continuous force regulation, no fatigue effect was observed highlighting the task-specific impact of fatigue. Similarly, Zabihhosseinian et al. (2020) demonstrated that cervical extensor fatigue reduced early tracing gains from ∼17% to ∼6%, again in a paradigm demanding fine visuomotor control.

**Table 2.**
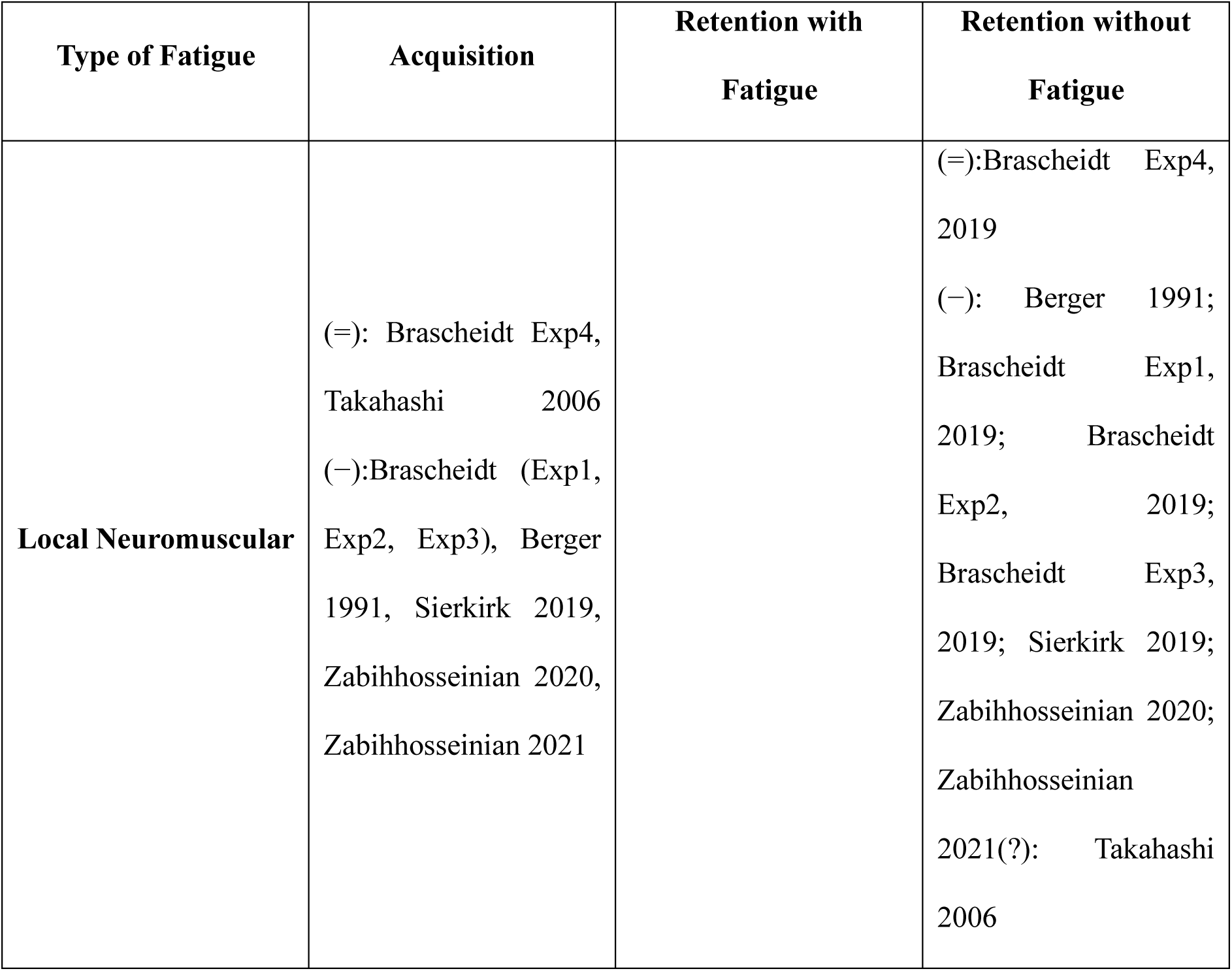
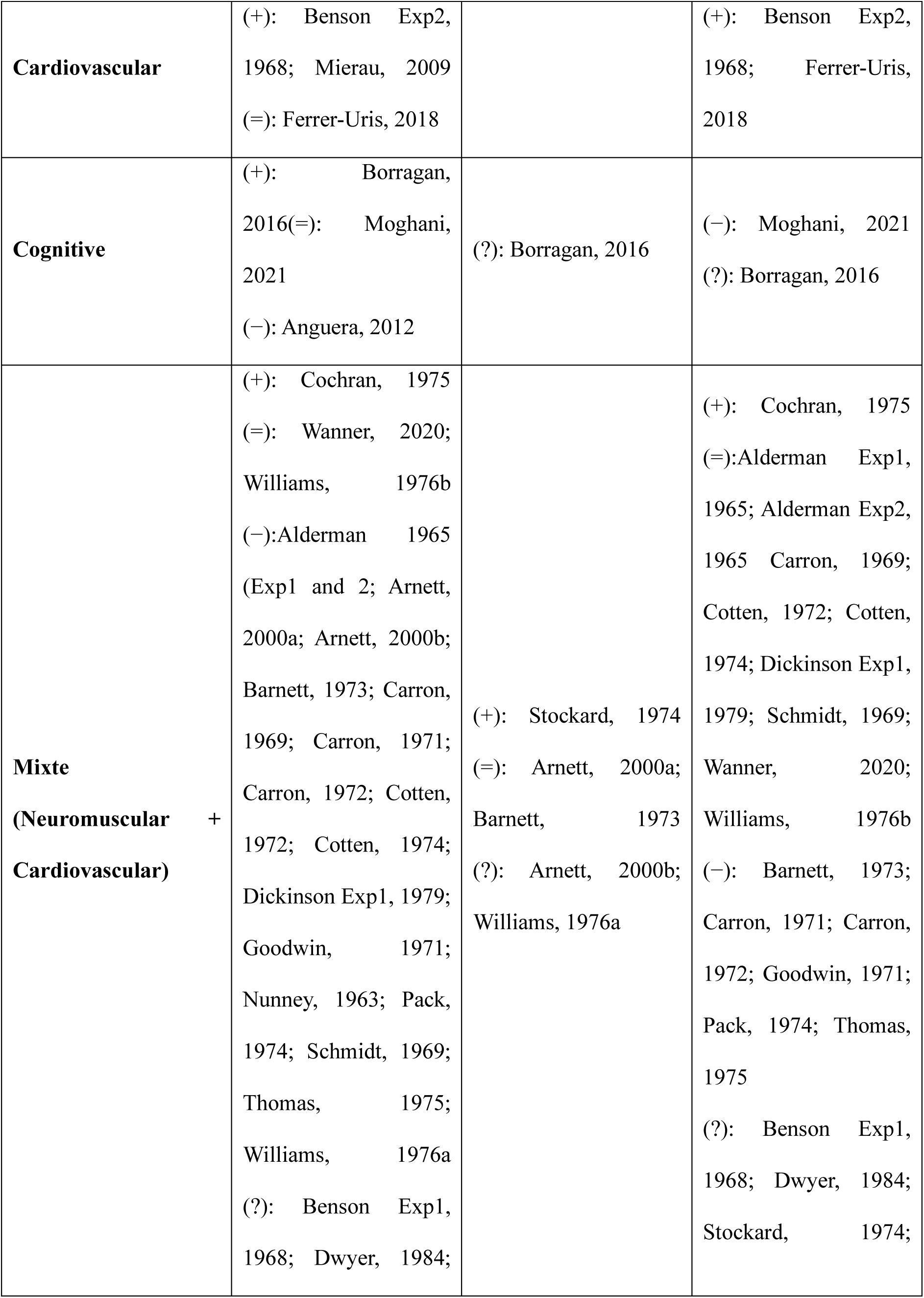
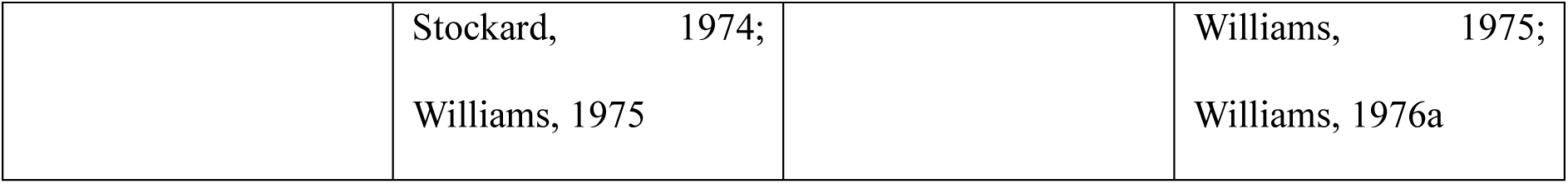
Fatigue Effects on Motor Learning: Evidence of Positive, Negative, and Neutral Outcomes. (+): Positive effect of fatigue on motor learning (fatigue improved performance); (−): Negative effect of fatigue (fatigue impaired performance); (=): Neutral effect (fatigue neither improved nor impaired performance); (?): Mixed or inconclusive results where effects were difficult to interpret.

Conversely, 10% of experiments (4/39) reported performance facilitation, mainly following systemic cardiovascular exertion (e.g., Mierau et al. (2009); Benson (1968), Exp. 2) or cognitively demanding protocols (Borragan et al., 2016). Neutral outcomes were observed in 15% of studies (6/39), whereas the remaining 10% (4/39) produced inconclusive results.

### 3.4. Effects of Fatigue on Retention After Recovery

Retention after recovery was evaluated in 34 of the 39 experiments (87%; see Table 2). Among these, 11 involved local neuromuscular fatigue: 3 experiments (27%) showed no retention decrement, 7 (64%) reported persistent impairments, and 1 (9%) was equivocal. Two studies assessed cardiovascular fatigue, and both demonstrated preserved or enhanced task performance retention. Cognitive fatigue was examined in two experiments: one (50%) found a clear retention deficit, while the other (50%) yielded inconclusive results. The largest subset, comprising 19 experiments, combined neuromuscular and cardiovascular protocols. Of these, 1 (5%) showed a facilitatory effect, 7 (37%) observed no significant effect, 6 (32%) reported lasting impairments, and 5 (26%) were equivocal.

### 3.5. Effects of Fatigue on Retention Under Sustained Fatigue

Retention with fatigue was assessed in 6 of the 39 experiments (15 %; see Table 2). No studies have assessed retention under isolated local neuromuscular or cardiovascular fatigue. One investigation of cognitive fatigue (Borragan et al., 2016) reported that retention benefited from a high-load session only when it was preceded by a low-load session, suggesting that transfer of adaptation depends on the sequence of cognitive demands. The remaining five studies combined neuromuscular and cardiovascular protocols, yielding mixed results. Notably, Arnett et al. (2000b)reported that men who trained while fatigued showed improved performance during a retention test, and they also performed with fatigue. However, women who trained with fatigue showed an impaired performance during a similar retention test. Furthermore, the only other experiment revealing an advantage of training with fatigue, when retention is also tested with fatigue, only included male participants (Stockard, 1974).

### 3.6. Neurophysiological Responses to Fatigue in Motor Learning

Four heterogeneous experiments incorporated included neurophysiological measures. In Branscheidt et al. (2019) Exp. 3, fatigue led to reduced MEP amplitudes, most likely reflecting a direct consequence of muscle fatigue rather than an interaction with sensorimotor learning. More importantly, when a depotentiation rTMS protocol was applied immediately after training, the typical retention deficits induced by fatigue were attenuated, suggesting that interference with plasticity mechanisms can mitigate the impact of fatigue on consolidation. Zabihhosseinian et al. (2020) found that cervical-extensor fatigue suppressed the decrease in cerebello-cortical inhibition, as measured with TMS, normally observed in their control group following a finger-tracing task. In a follow-up study, Zabihhosseinian et al. (2021) reported that the same fatigue protocol increased N24 and N30 somatosensory-evoked potentials after motor training, while the N24 component decreased in controls. In both studies, fatigue was associated with smaller acquisition and retention gains. Finally, Mierau et al. (2009) observed that exhausting running before a visuomotor inversion task improved tracing performance and was accompanied by reduced beta-band power but unchanged alpha-band power within the EEG oscillations compared to the rested condition.

## 4. Discussion

This systematic review assessed how neuromuscular, cardiovascular and cognitive fatigue influence motor learning. Across thirty-nine experiments, local neuromuscular fatigue generally hampered both acquisition and retention. By contrast, acute systemic cardiovascular fatigue tended to enhance task acquisition and retention, although this effect has been less extensively studied (Benson, 1968; Ferrer-Uris et al., 2018; Mierau et al., 2009; Neva et al., 2019). Mixed protocols that combined neuromuscular and cardiovascular strain consistently slowed initial acquisition, yet retention outcomes ranged from preserved to impaired, a variability potentially shaped by task demands, testing state and participant characteristics (Arnett et al., 2000a; Barnett et al., 1973; Carron, 1969; Cochran, 1975; Godwin & Schmidt, 1971). The few studies that evaluated the effects of cognitive fatigue on motor learning were equivocal both during the acquisition and retention tests. (Anguera et al., 2012; Borragan et al., 2016; Khojasteh Moghani et al., 2021). Across all types of fatigue, very few studies evaluated retention when participants were fatigued again. Some of these experiments showed continuing impairment and others reported preserved performance or even enhanced performance for participants who trained while experiencing fatigue, underscoring the context dependence of fatigue effects across early acquisition and subsequent recall.

### 4.1. Neuromuscular fatigue

Local neuromuscular fatigue may interfere with motorng learning through converging peripheral and central mechanisms, most of which are primarily documented in the context of motor control. Peripherally, metabolite accumulation and enhanced group III–IV afferent activity degrade proprioceptive accuracy, increase noise within the sensorimotor loop, and reduce the reliability of sensory feedback used for error correction and adaptation (Franklin & Wolpert, 2011; Shadmehr & Krakauer, 2008). Importantly, fatigue does not necessarily suppress plasticity but may bias it. demonstrated that muscle fatigue impaired sequence learning and reduced motor-evoked potentials. Strikingly, applying a depotentiation rTMS protocol after practice attenuated the usual retention deficits, suggesting that maladaptive plasticity had been consolidated under fatigue, and that disrupting this consolidation reduced the impairment. Supporting this view, Zabihhosseinian et al. (2020, 2021) reported that cervical muscle fatigue modulated cerebellar-corticalinhibition and amplified early somatosensory-evoked potentials, suggesting altered proprioceptive integration and reduced fidelity of sensorimotor representations. Similarly,(Nicholas et al., 2019) observed that limb-specific fatigue impaired both acquisition and retention of a proprioceptive task, further emphasizing the role of sensory degradation. Adding nuance, Pawłowski et al. (2024) showed that fatigue induces subtle reorganizations of motor synergies at different levels of neuromotor control while global task performance remains stable. This finding supports the notion that fatigue reshapes motor coordination strategies rather than producing a uniform decline. Taken together, these findings suggest that neuromuscular fatigue does not merely reduce plasticity but can bias it towards alternative coordination patterns. Whether these patterns are maladaptive or adaptive may depend on the testing context: they appear suboptimal when retention is assessed in a rested state, but could be advantageous when performance is recalled under similar fatigue conditions. In this sense, fatigue imposes a dual constraint on learning: it lowers excitability while shaping motor representations in a context-dependent manner.

### 4.2. Cardiovascular fatigue

Vigorous aerobic exercise often acts as a short-lived primer of plasticity rather than a constraint on learning, a phenomenon supported by both experimental and meta-analytic evidence (Dal Maso et al., 2018; Wanner et al., 2020). After brief recovery, circulating catecholamines and neurotrophic factors such as dopamine, norepinephrine, and BDNF increase, cortical excitability rises, and the probability of synaptic strengthening is enhanced, producing a transient learning-ready state sometimes termed exercise-induced priming (Ferrer-Uris et al., 2018; Mierau et al., 2009; Neva et al., 2019). Classic and contemporary studies indicate that exercise performed shortly before or immediately after practice improves retention, even when immediate acquisition remains unchanged, in sharp contrast to the noise-inducing effects of local muscular fatigue (Benson, 1968; Mierau et al., 2009). Evidence for peripheral effects of cardiovascular exercise on motor learning remains limited but informative. A single bout of aerobic exercise has been reported to transiently improve joint position sense and postural balance, suggesting a short-term facilitation of proprioceptive acuity and sensorimotor control (Harry-Leite et al., 2022). In contrast, when exercise is exhaustive, maximal voluntary strength can be transiently depressed due to peripheral fatigue (Lebesque et al., 2022). However, under moderate aerobic loads, voluntary force production appears to be preserved, indicating that cardiovascular exercise at tolerable intensity does not compromise basic motor output and may even support stable force control (Slimani et al., 2016)

### 4.3. Mixed fatigue

When neuromuscular and cardiovascular components are combined, outcomes vary substantially across tasks and testing states. During acquisition, decrements are frequently observed, with slower learning rates and higher error scores when fatigue is introduced during practice or immediately before training particularly in protocols that combine localized muscular effort with aerobic exertion (Carron, 1969; Cotten et al., 1972; Dickinson et al., 1979). These impairments are often attributed to the predominance of peripheral mechanisms, as neuromuscular fatigue reduces the accuracy of proprioceptive input and compromises force precision, thereby limiting the effectiveness of error correction processes during early learning.In contrast, the priming effects of cardiovascular exertion are more likely to emerge after partial recovery rather than during the acquisition phase. Retention results remain heterogeneous. Several studies reported that when participants were retested in a fresh state, performance between fatigued and non-fatigued groups converged, consistent with the idea that offline consolidation can partly compensate for early deficits (Arnett et al., 2000a; Barnett et al., 1973). However, other experiments observed persistent impairments despite recovery, indicating that consolidation does not uniformly offset the negative effects of fatigue (Carron, 1969; Carron, 1972; Godwin & Schmidt, 1971)Thus, while most studies suggest a neutral impact of mixed fatigue on retention in a rested state, the evidence remains too inconsistent to draw firm conclusions. Even less is known about retention under renewed fatigue. The very few available experiments point in different directions, ranging from state-dependent advantages where participants who practiced while fatigued performed better when tested again in a fatigued state (Stockard, 1974; Williams & Cooper, 1976) to continued performance decrements. Given the small number and methodological variability of these studies, it remains unclear whether training under fatigue provides any reliable state-dependent benefits for recall.

### 4.4. Methodological strengths and limitations

This review drew on a multi-database search, applied a uniform task taxonomy, and implemented duplicate screening and data extraction, which increases reliability. Study quality was appraised with the Downs and Black checklist, and methodological rigor improved across publication years, although heterogeneity remains evident. Several constraints limit inference. Fatigue protocols varied widely in modality, intensity, and timing, and motor tasks ranged from simple tracking to complex sequence learning, which precluded formal meta-analysis and complicated direct comparisons. Follow-up rarely exceeded 48 hours, leaving the durability of fatigue effects on motor learning underexplored. Only a handful of experiments incorporated neurophysiological measures, which limits mechanistic conclusions. Very few studies assessed retention in the same fatigued state as acquisition, preventing firm conclusions about whether neuromuscular fatigue continues to depress performance when both learning and recall occur under fatigue. Individual moderators, such as sex distribution, baseline fitness, or skill level, were seldom analyzed, despite preliminary evidence suggesting that they may shape responses to fatigue. For example, studies that stratified by sex or training status hinted at differential effects but did not consistently test these interactions (Arnett et al., 2000b; Barnett et al., 1973; Wanner et al., 2020).

### 4.5. Practical implications

Technical drills should ideally be scheduled when neuromuscular fatigue is minimal, since proprioceptive accuracy and corticospinal excitability both critical for stable encoding are reduced under fatigue (Branscheidt et al., 2019; Zabihhosseinian et al., 2020; Zabihhosseinian et al., 2021)If skills are expected to be performed in a fatigued state, however, the principle of practice specificity suggests that training under fatigue could facilitate state-dependent recall (Arnett et al., 2000b), though such benefits may come at the cost of weaker transfer to fresh conditions. Moderate aerobic exercise followed by short recovery may serve as a plasticity primer, enhancing consolidation when practice occurs after acute sensations of exhaustion have subsided (Ferrer-Uris et al., 2018; Mierau et al., 2009). Proprioceptive “refreshers,” such as light balance tasks, might help restore accuracy before resuming skill practice, but this remains hypothetical and requires validation in motor learning contexts (Proske & Gandevia, 2012). Likewise, neuromodulation strategies such as facilitatory rTMS may counteract fatigue-induced reductions in excitability, though their applicability in training settings remains to be demonstrated. Ultimately, implementation should be individualized: coaches and clinicians should monitor both subjective and objective fatigue markers and tailor workloads to the athlete’s or patient’s profile.

### 4.6. Future research directions

Future work should address three priorities. First, retention should be tested both in fresh and fatigued states to clarify whether state-dependent learning occurs. Second, studies should incorporate neurophysiological measures (e.g., TMS, EEG) to link excitability, inhibition, and sensory processing with learning outcomes. Third, randomized trials should evaluate countermeasures such as aerobic priming, neuromodulation, or recovery strategies to establish evidence-based approaches for optimizing skill acquisition under fatigue. Expanding samples beyond young healthy adults, including older or clinical populations, would also improve external validity.

## 5. Conclusion

This systematic review highlights the complex and context-dependent influence of fatigue on motor learning. Local neuromuscular fatigue consistently hampers acquisition and often reduces retention, primarily by degrading proprioceptive accuracy and corticospinal excitability, although some evidence suggests that altered coordination patterns may be reinforced when learning occurs under fatigue. By contrast, cardiovascular exertion, when followed by short recovery, appears to act as a transient primer of neuroplasticity, improving consolidation and retention without compromising immediate performance. The effects of cognitive fatigue remain equivocal, with preliminary evidence indicating potential impairments in retention under high mental load but insufficient data for firm conclusions. Importantly, very few studies have assessed retention under fatigue, leaving open the question of whether state-dependent learning confers advantages when skills must later be performed in fatigued contexts. Methodological heterogeneity, short retention intervals, and limited neurophysiological measures further constrain interpretation. Overall, the evidence suggests that fatigue does not uniformly impair motor learning but instead modulates plasticity in ways that depend on the type of fatigue, the timing of assessment, and the nature of the task. Future studies should extend retention testing, integrate mechanistic measures, and explore countermeasures such as aerobic priming or neuromodulation to establish evidence-based strategies for optimizing training and rehabilitation when fatigue is unavoidable.

## Declaration of Generative AI

During the preparation of this work, the authors used Microsoft Copilot to check grammar and improve the clarity of the text. All suggestions were reviewed and edited by the authors, who take full responsibility for the final content of the article.

## Annexe

**Supplementary Table 1:**
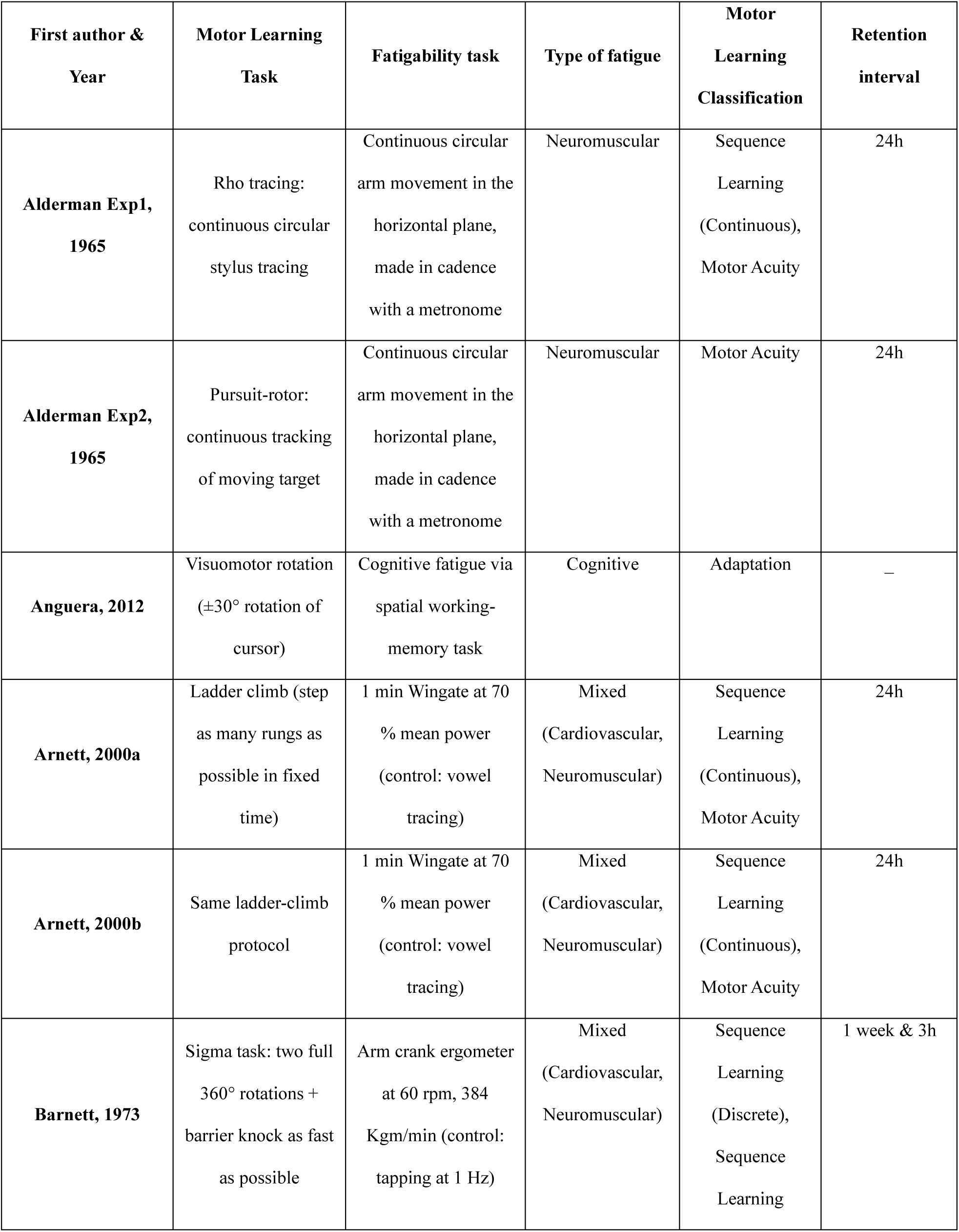
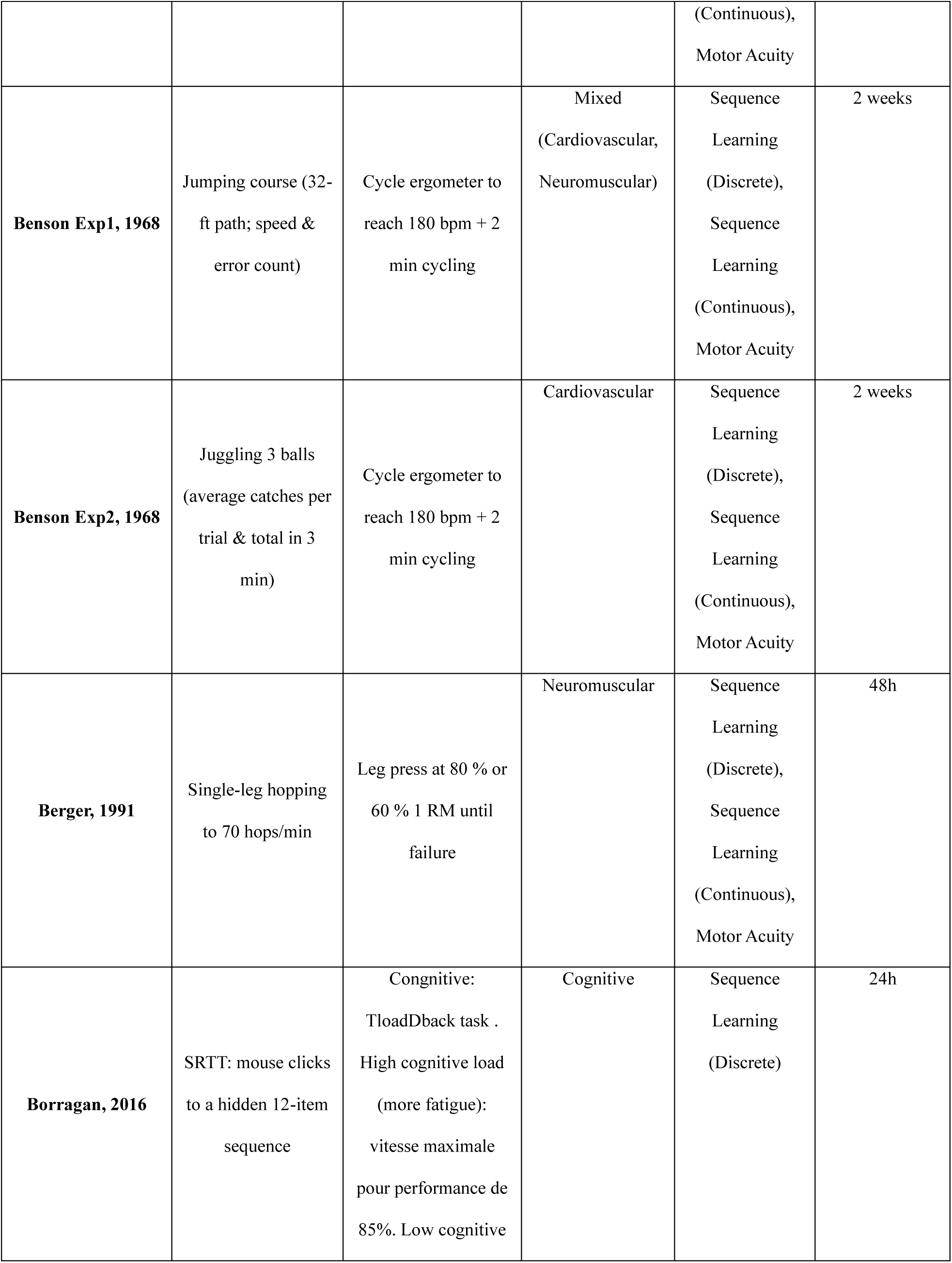
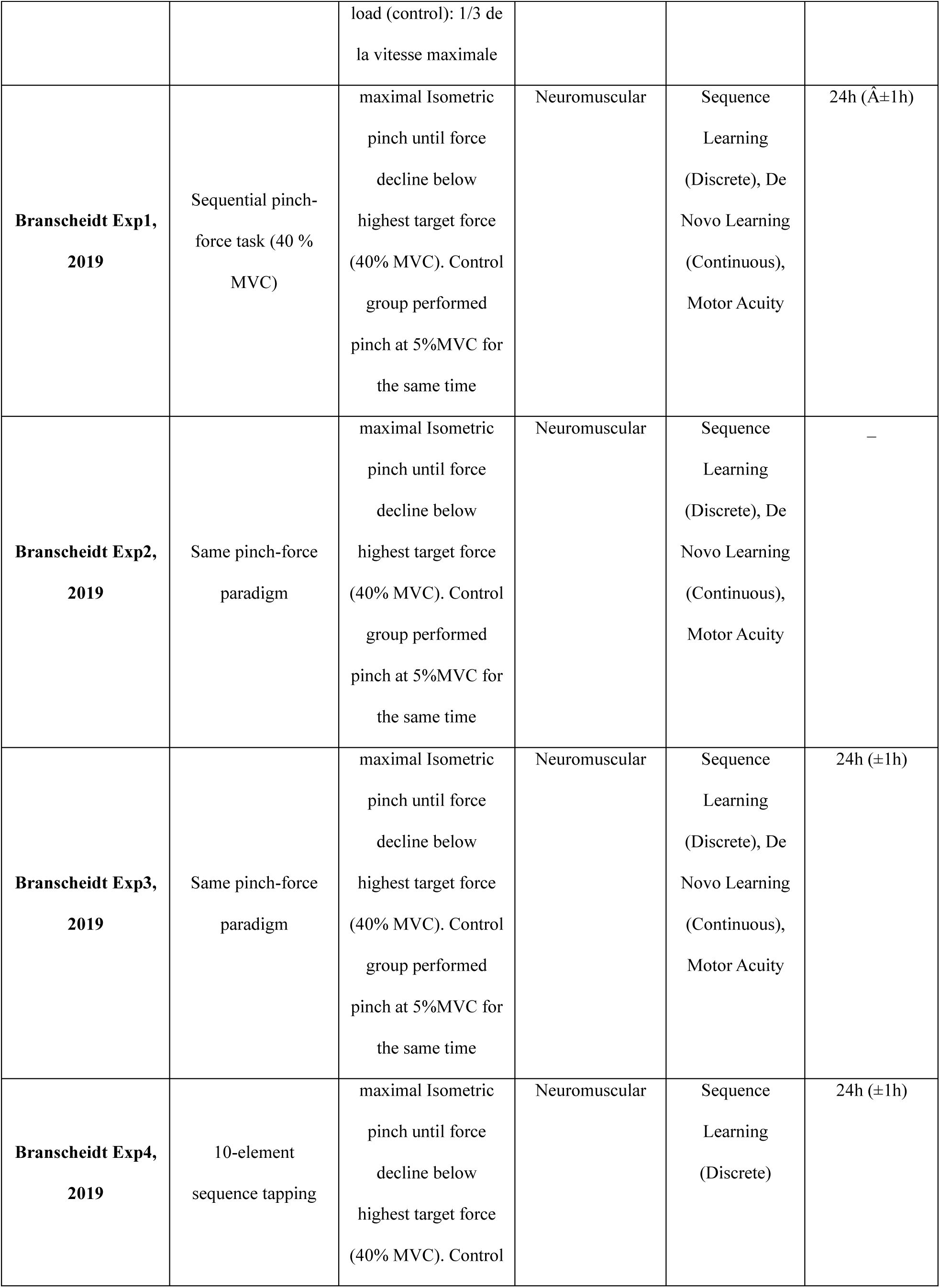
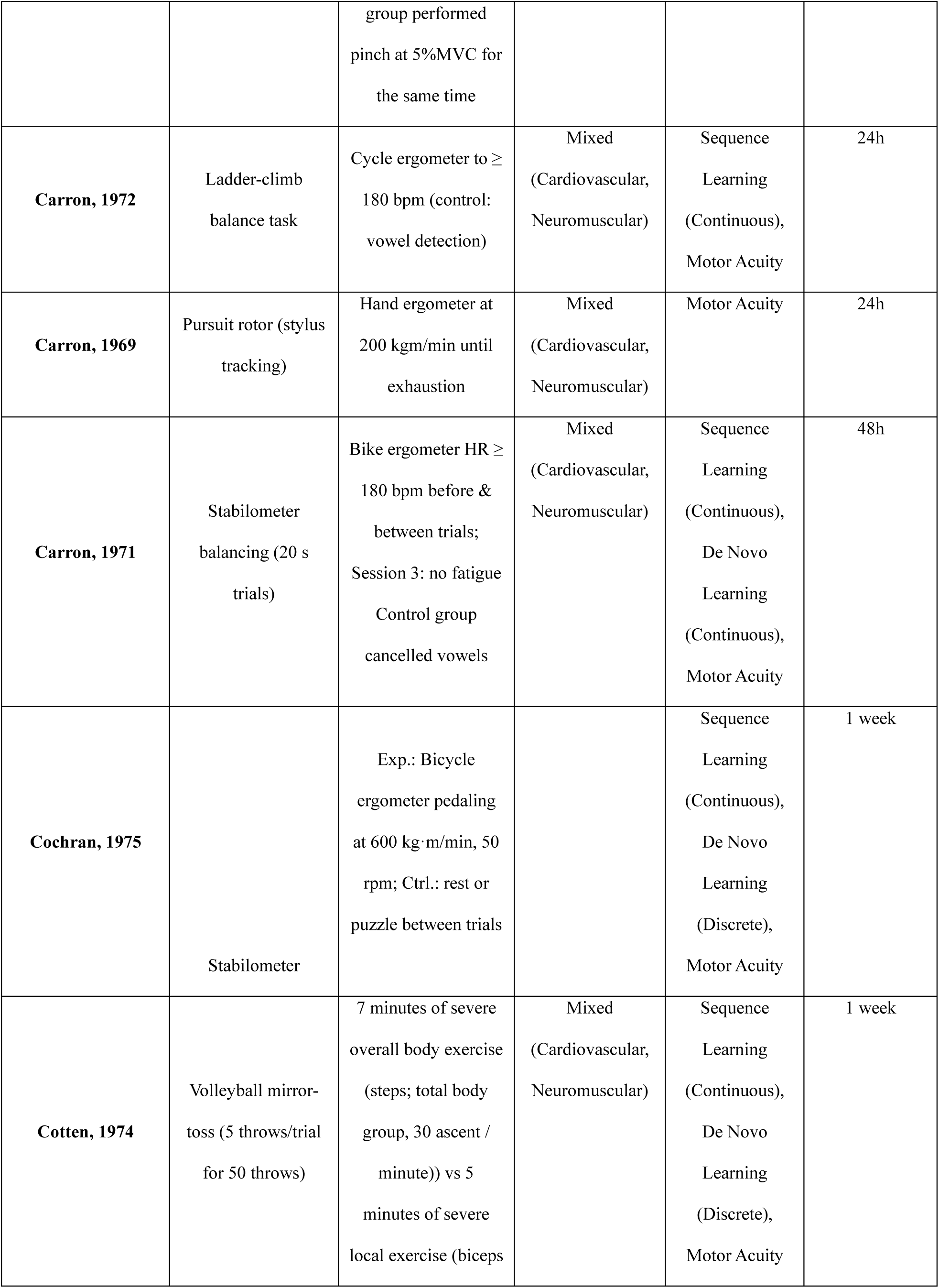
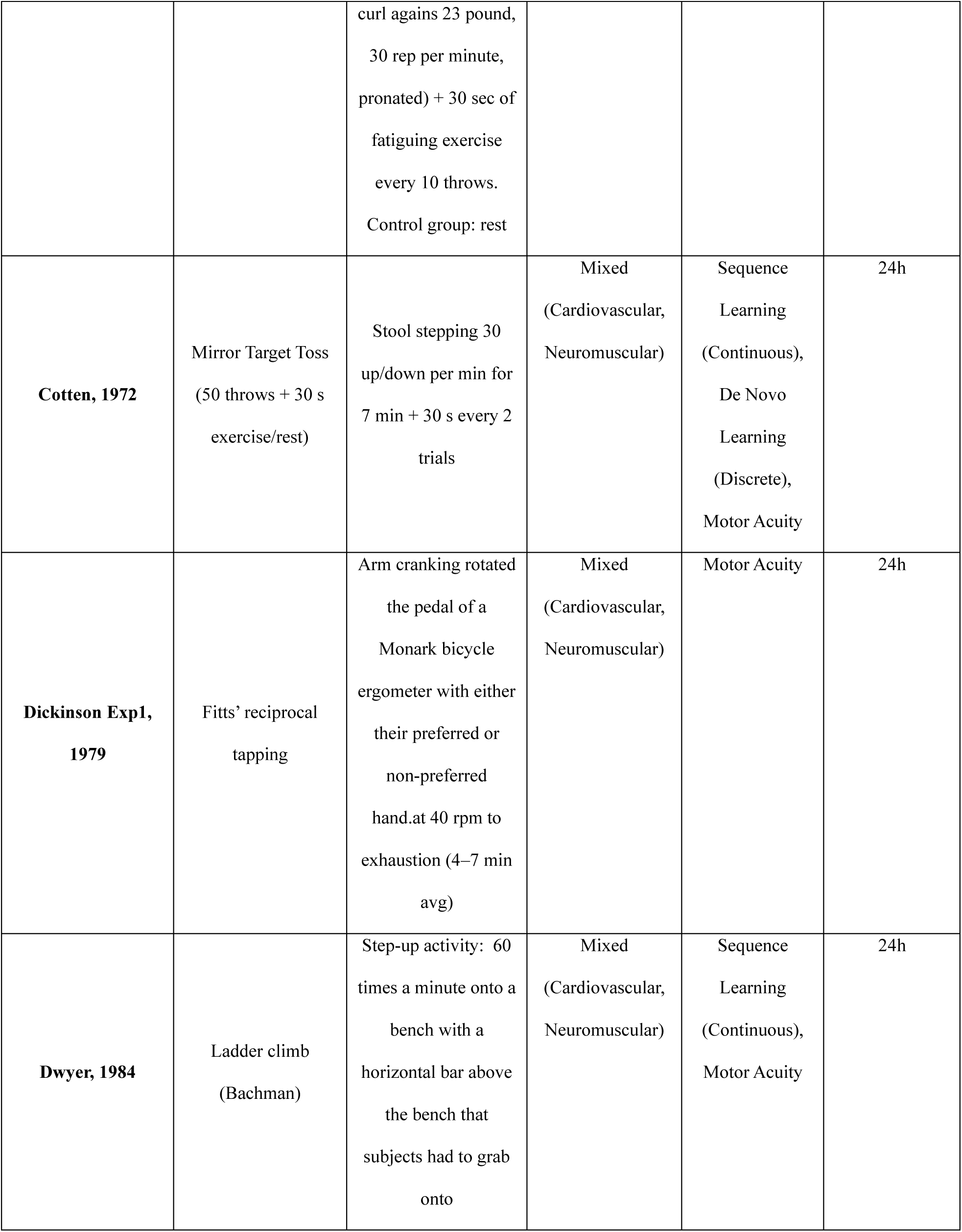
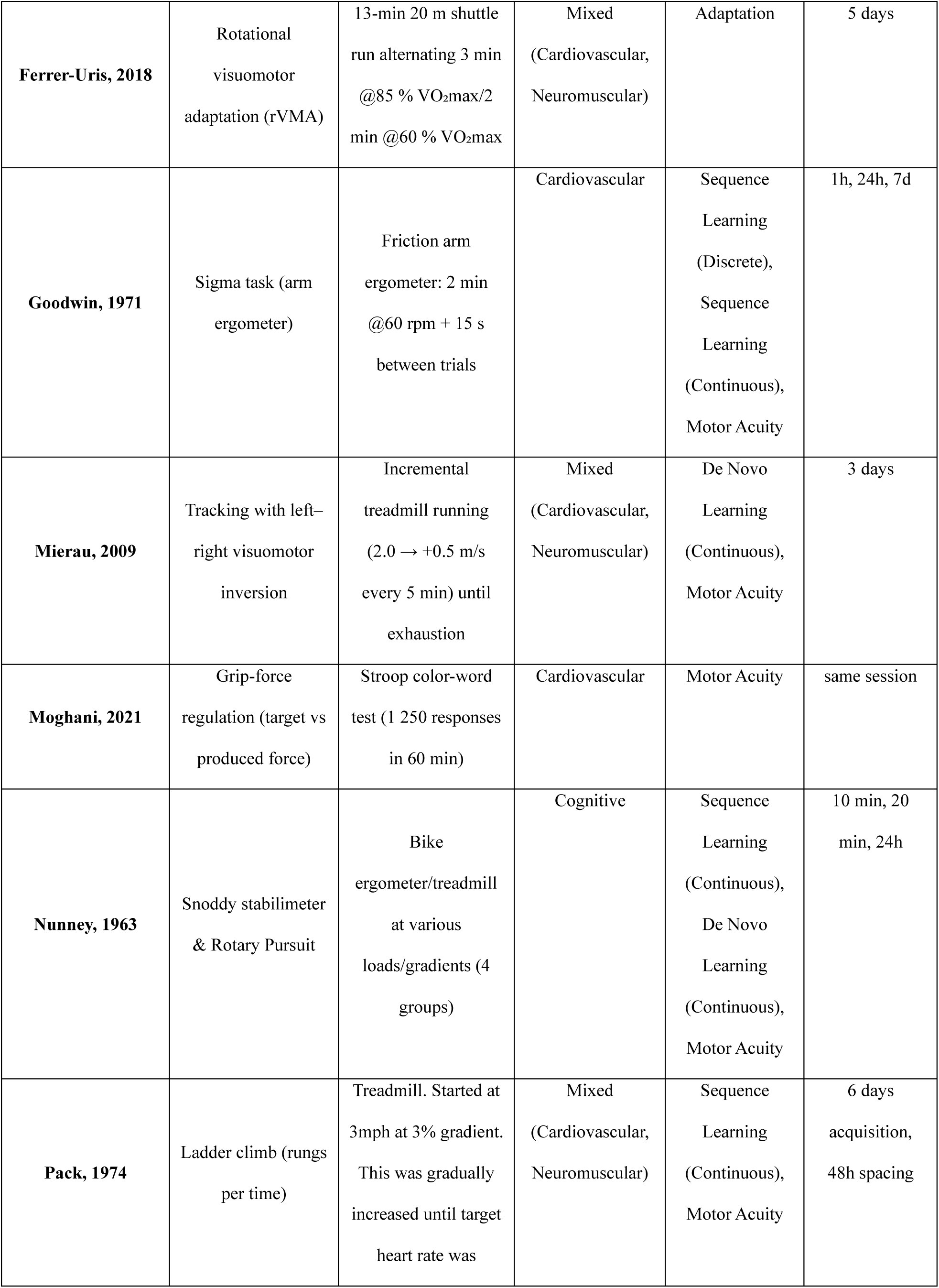
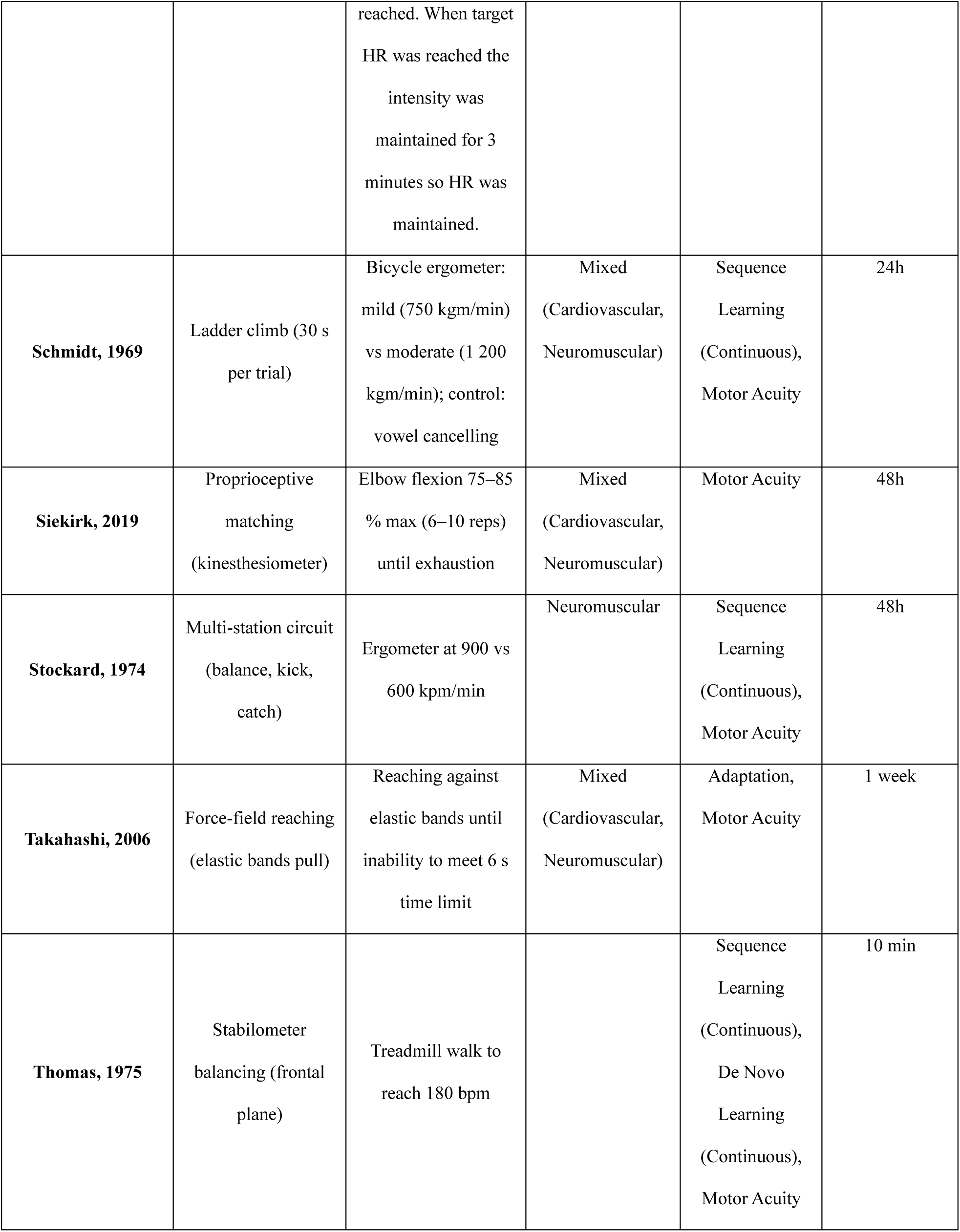
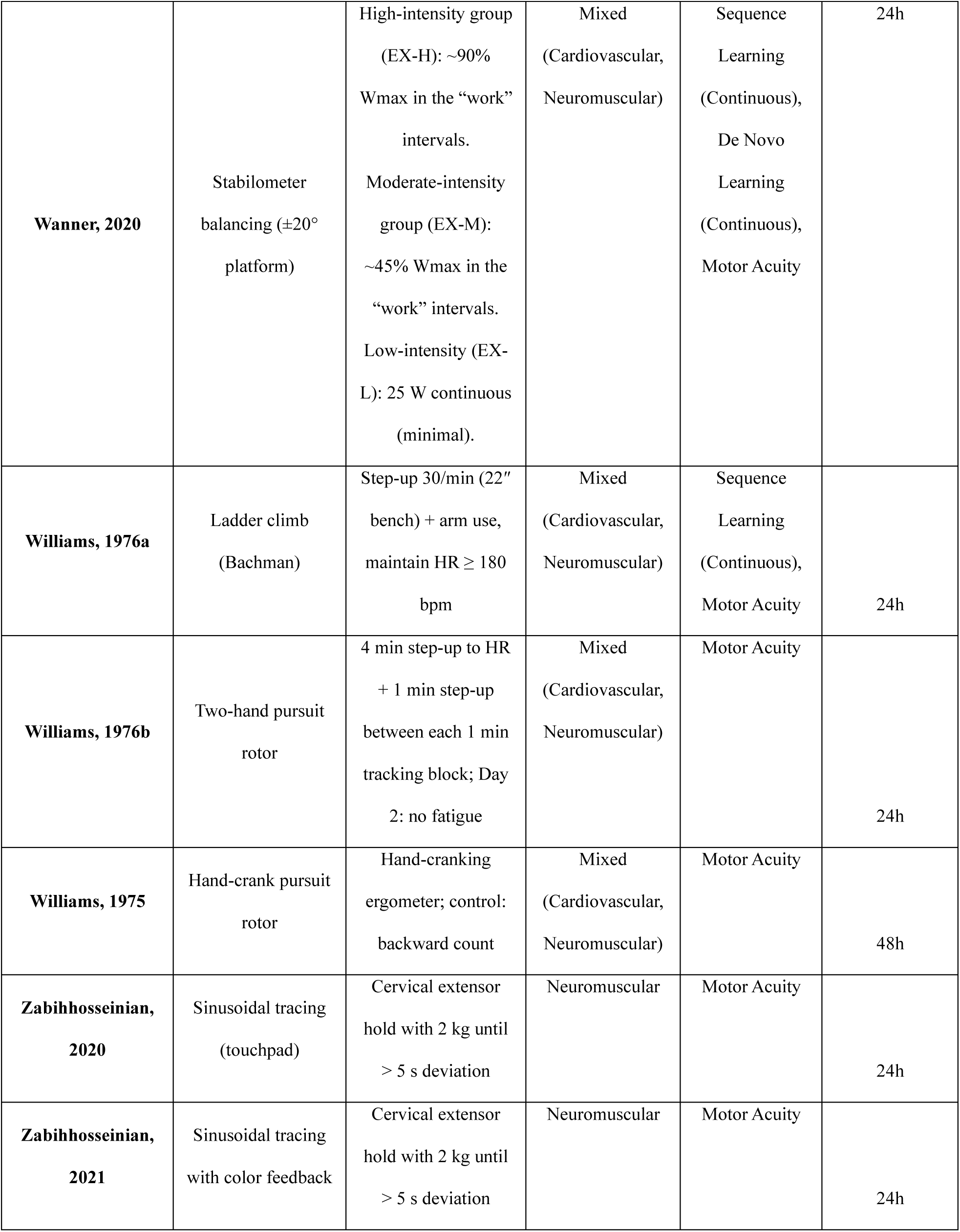
Fatigue and motor learning classification.

## References

Anguera, J. A., Bernard, J. A., Jaeggi, S. M., Buschkuehl, M., Benson, B. L., Jennett, S., Humfleet, J., Reuter-Lorenz, P. A., Jonides, J., & Seidler, R. D. (2012). The effects of working memory resource depletion and training on sensorimotor adaptation. Behavioural Brain Research, 228(1), 107–115.

Arnett, DeLuccia, D., & Gilmartin, K. (2000b). Male and female differences and the specificity of fatigue on skill acquisition and transfer performance. Res Q Exerc Sport, 71(2), 201–205. 10.1080/02701367.2000.10608899

Arnett, M. G., Bennett, C., Gilmartin, K., & DeLuccia, D. (2000a). The Effect of Specific Anaerobically Induced Fatigue on Skill Acquisition and Transfer Performance. Physical Educator, 57(4).

Barnett, M. L., Ross, D., Schmidt, R. A., & Todd, B. (1973). Motor Skills Learning and the Specificity of Training Principle. Research Quarterly. American Association for Health, Physical Education and Recreation, 44(4), 440–447. 10.1080/10671188.1973.10615224

Benson, D. W. (1968). Influence of imposed fatigue on learning a jumping task and a juggling task. Research quarterly, 39(2), 251–257.

Borragan, G., Slama, H., Destrebecqz, A., & Peigneux, P. (2016). Cognitive Fatigue Facilitates Procedural Sequence Learning. Front Hum Neurosci, 10, 86. 10.3389/fnhum.2016.00086

Branscheidt, M., Kassavetis, P., Anaya, M., Rogers, D., Huang, H. D., Lindquist, M. A., & Celnik, P. (2019). Fatigue induces long-lasting detrimental changes in motor-skill learning. Elife, 8. 10.7554/eLife.40578

Carron, A. V. (1969). Physical fatigue and motor learning. Research quarterly, 40(4), 682–686.

Carron, A. V. (1972). Motor performance and learning under physical fatigue. Medicine & Science in Sports & Exercise, 4(2), 101-106. 10.1249/00005768-197200420-00010

Cochran, B. J. (1975). Effect of physical fatigue on learning to perform a novel motor task. Research quarterly, 46(2), 243–249.

Cotten, D. J., Thomas, J. R., Spieth, W. R., & Biasiotto, J. (1972). Temporary fatigue effects in a gross motor skill. J Mot Behav, 4(4), 217–222. 10.1080/00222895.1972.10734937

Dal Maso, F., Desormeau, B., Boudrias, M.-H., & Roig, M. (2018). Acute cardiovascular exercise promotes functional changes in cortico-motor networks during the early stages of motor memory consolidation. NeuroImage, 174, 380–392.

Davranche, K., Brisswalter, J., & Radel, R. (2015). Where are the limits of the effects of exercise intensity on cognitive control? Journal of Sport and Health Science, 4(1), 56–63.

Dickinson, J., Medhurst, C., & Whittingham, N. (1979). Warm-and fatigue in skill acquisition and performance. J Mot Behav, 11(1), 81–86. 10.1080/00222895.1979.10735175

Downs, S. H., & Black, N. (1998). The feasibility of creating a checklist for the assessment of the methodological quality both of randomised and non-randomised studies of health care interventions. Journal of epidemiology & community health, 52(6), 377–384.

Enoka, R. M., & Duchateau, J. (2008). Muscle fatigue: what, why and how it influences muscle function. The Journal of physiology, 586(1), 11–23.

Enoka, R. M., & Duchateau, J. (2016). Translating fatigue to human performance. Medicine and science in sports and exercise, 48(11), 2228.

Ferrer-Uris, B., Busquets, A., & Angulo-Barroso, R. (2018). Adaptation and Retention of a Perceptual-Motor Task in Children: Effects of a Single Bout of Intense Endurance Exercise. J Sport Exerc Psychol, 40(1), 1–9. 10.1123/jsep.2017-0044

Franklin, D. W., & Wolpert, D. M. (2011). Computational mechanisms of sensorimotor control. Neuron, 72(3), 425–442.

Godwin, M. A., & Schmidt, R. A. (1971). Muscular Fatigue and Learning a Discrete Motor Skill. Research Quarterly. American Association for Health, Physical Education and Recreation, 42(4), 374–382. 10.1080/10671188.1971.10615084

Harry-Leite, P., Paquete, M., Teixeira, J., Santos, M., Sousa, J., Fraiz-Brea, J. A., & Ribeiro, F. (2022). Acute impact of proprioceptive exercise on proprioception and balance in athletes. Applied Sciences, 12(2), 830.

Khojasteh Moghani, M., Zeidabadi, R., Shahabi Kaseb, M. R., & Bahreini Borujeni, I. (2021). Mental Fatigue Reduces the Benefits of Self-Controlled Feedback on Learning a Force Production Task. Percept Mot Skills, 128(5), 2398–2414. 10.1177/00315125211037306

Krakauer, J. W., Hadjiosif, A. M., Xu, J., Wong, A. L., & Haith, A. M. (2019). Motor Learning. Compr Physiol, 9(2), 613–663. 10.1002/cphy.c170043

Kwon, Y. H., Kwon, J. W., & Lee, M. H. (2015). Effectiveness of motor sequential learning according to practice schedules in healthy adults; distributed practice versus massed practice. Journal of physical therapy science, 27(3), 769–772.

Lebesque, L., Scaglioni, G., & Martin, A. (2022). The impact of submaximal fatiguing exercises on the ability to generate and sustain the maximal voluntary contraction. Frontiers in physiology, 13, 970917. 10.3389/fphys.2022.970917

Lehmann, N., Villringer, A., & Taubert, M. (2022). Priming cardiovascular exercise improves complex motor skill learning by affecting the trajectory of learning-related brain plasticity. Scientific reports, 12(1). 10.1038/s41598-022-05145-7

Marcora, S. M., Staiano, W., & Manning, V. (2009). Mental fatigue impairs physical performance in humans. Journal of applied physiology, 106(3), 857–864.

Marin Bosch, B., Bringard, A., Logrieco, M. G., Lauer, E., Imobersteg, N., Thomas, A., Ferretti, G., Schwartz, S., & Igloi, K. (2020). Effect of acute physical exercise on motor sequence memory. Scientific reports, 10(1), 15322.

Mierau, A., Schneider, S., Abel, T., Askew, C., Werner, S., & Struder, H. K. (2009). Improved sensorimotor adaptation after exhaustive exercise is accompanied by altered brain activity. Physiol Behav, 96(1), 115–121. 10.1016/j.physbeh.2008.09.002

Moriarty, T., Johnson, A., Thomas, M., Evers, C., Auten, A., Cavey, K., Dorman, K., & Bourbeau, K. (2022). Acute Aerobic Exercise-Induced Motor Priming Improves Piano Performance and Alters Motor Cortex Activation [Original Research]. Frontiers in Psychology, Volume 13 - 2022. 10.3389/fpsyg.2022.825322

Neva, J. L., Ma, J. A., Orsholits, D., Boisgontier, M. P., & Boyd, L. A. (2019). The effects of acute exercise on visuomotor adaptation, learning, and inter-limb transfer. Experimental brain research, 237(4), 1109–1127. 10.1007/s00221-019-05491-5

Nicholas, S., Qin, L., & Bradley, K. (2019). Effects of Limb-Specific Fatigue on Motor Learning during an Upper Extremity Proprioceptive Task. International Journal of Motor Control and Learning, 1(2), 41–46.

Pageaux, B., & Lepers, R. (2018). The effects of mental fatigue on sport-related performance. Progress in brain research, 240, 291–315.

Pawłowski, M., Furmanek, M. P., & Juras, G. (2024). Does muscle fatigue change motor synergies at different levels of neuromotor control? Frontiers in Human Neuroscience, 18, 1519462. 10.3389/fnhum.2024.1519462

Proske, U., & Gandevia, S. C. (2012). The proprioceptive senses: their roles in signaling body shape, body position and movement, and muscle force. Physiological reviews.

Russell, M. S., Vasilounis, S. S., Lefebvre, E., Drake, J. D. M., & Chopp-Hurley, J. N. (2024). Variability in musculoskeletal fatigue responses associated with repeated exposure to an occupational overhead drilling task completed on successive days. Human Movement Science, 97. 10.1016/j.humov.2024.103276

Salihu, A. T., Hill, K. D., & Jaberzadeh, S. (2022). Effect of cognitive task complexity on dual task postural stability: a systematic review and meta-analysis. Experimental brain research, 240(3), 703–731.

Shadmehr, R., & Krakauer, J. W. (2008). A computational neuroanatomy for motor control. Exp Brain Res, 185(3), 359–381. 10.1007/s00221-008-1280-5

Slimani, M., Chamari, K., Miarka, B., Del Vecchio, F. B., & Cheour, F. (2016). Effects of plyometric training on physical fitness in team sport athletes: a systematic review. Journal of human kinetics, 53, 231.

Stockard, J. R. (1974). Prior Physical Exertion in Learning a Novel Gross-Motor Task. Perceptual and Motor Skills, 38(1), 146. 10.2466/pms.1974.38.1.146

Takahashi, C. D., Nemet, D., Rose-Gottron, C. M., Larson, J. K., Cooper, D. M., & Reinkensmeyer, D. J. (2006). Effect of muscle fatigue on internal model formation and retention during reaching with the arm. J Appl Physiol (1985), 100(2), 695-706. 10.1152/japplphysiol.00140.2005

Taylor, J. L. (2018). Motor Control And Motor Learning Under Fatigue Conditions. In.

Taylor, J. L., & Gandevia, S. C. (2008). A comparison of central aspects of fatigue in submaximal and maximal voluntary contractions. Journal of applied physiology, 104(2), 542–550.

Wanner, P., Muller, T., Cristini, J., Pfeifer, K., & Steib, S. (2020). Exercise Intensity Does not Modulate the Effect of Acute Exercise on Learning a Complex Whole-Body Task. Neuroscience, 426, 115–128. 10.1016/j.neuroscience.2019.11.027

Williams, L. R. T., & Cooper, E. L. (1976). Fatigue in Learning and Performance of a Gross Tracking Task. Perceptual and Motor Skills, 42, 1287–1294. 10.2466/pms.1976.42.3c.1287

Youssef, L., Harroum, N., Francisco, B. A., Johnson, L., Arvisais, D., Pageaux, B., Romain, A. J., Hayward, K. S., & Neva, J. L. (2024). Neurophysiological effects of acute aerobic exercise in young adults: a systematic review and meta-analysis. Neuroscience & Biobehavioral Reviews, 164, 105811.

Zabihhosseinian, M., Yielder, P., Berkers, V., Ambalavanar, U., Holmes, M., & Murphy, B. (2020). Neck muscle fatigue impacts plasticity and sensorimotor integration in cerebellum and motor cortex in response to novel motor skill acquisition. J Neurophysiol, 124(3), 844–855. 10.1152/jn.00437.2020

Zabihhosseinian, M., Yielder, P., Wise, R., Holmes, M., & Murphy, B. (2021). Effect of Neck Muscle Fatigue on Hand Muscle Motor Performance and Early Somatosensory Evoked Potentials. Brain Sci, 11(11). 10.3390/brainsci11111481

