## Supplementary Table 1 for "Neuromuscular, Cardiovascular, and Cognitive Fatigue in Motor Learning: A Systematic Review"

| **First author & Year** | **Motor Learning Task** | **Fatigability task** | **Type of fatigue** | **Motor Learning Classification** | **Retention interval** |
|---|---|---|---|---|---|
| **Alderman Exp1, 1965** | Rho tracing: continuous circular stylus tracing | Continuous circular arm movement in the horizontal plane, made in cadence with a metronome | Neuromuscular | Sequence Learning (Continuous), Motor Acuity | 24h |
| **Alderman Exp2, 1965** | Pursuit-rotor: continuous tracking of moving target | Continuous circular arm movement in the horizontal plane, made in cadence with a metronome | Neuromuscular | Motor Acuity | 24h |
| **Anguera, 2012** | Visuomotor rotation (±30° rotation of cursor) | Cognitive fatigue via spatial working-memory task | Cognitive | Adaptation | _ |
| **Arnett, 2000a** | Ladder climb (step as many rungs as possible in fixed time) | 1 min Wingate at 70 % mean power (control: vowel tracing) | Mixed (Cardiovascular, Neuromuscular) | Sequence Learning (Continuous), Motor Acuity | 24h |
| **Arnett, 2000b** | Same ladder-climb protocol | 1 min Wingate at 70 % mean power (control: vowel tracing) | Mixed (Cardiovascular, Neuromuscular) | Sequence Learning (Continuous), Motor Acuity | 24h |
| **Barnett, 1973** | Sigma task: two full 360° rotations + barrier knock as fast as possible | Arm crank ergometer at 60 rpm, 384 Kgm/min (control: tapping at 1 Hz) | Mixed (Cardiovascular, Neuromuscular) | Sequence Learning (Discrete), Sequence Learning (Continuous), Motor Acuity | 1 week & 3h |

| | | | | | |
|---|---|---|---|---|---|
| **Benson Exp1, 1968** | Jumping course (32-ft path; speed & error count) | Cycle ergometer to reach 180 bpm + 2 min cycling | Mixed (Cardiovascular, Neuromuscular) | Sequence Learning (Discrete), Sequence Learning (Continuous), Motor Acuity | 2 weeks |
| **Benson Exp2, 1968** | Juggling 3 balls (average catches per trial & total in 3 min) | Cycle ergometer to reach 180 bpm + 2 min cycling | Cardiovascular | Sequence Learning (Discrete), Sequence Learning (Continuous), Motor Acuity | 2 weeks |
| **Berger, 1991** | Single-leg hopping to 70 hops/min | Leg press at 80 % or 60 % 1 RM until failure | Neuromuscular | Sequence Learning (Discrete), Sequence Learning (Continuous), Motor Acuity | 48h |
| **Borragan, 2016** | SRTT: mouse clicks to a hidden 12-item sequence | Congnitive: TloadDback task . High cognitive load (more fatigue): vitesse maximale pour performance de 85%. Low cognitive load (control): 1/3 de la vitesse maximale | Cognitive | Sequence Learning (Discrete) | 24h |

| | | | | | |
|---|---|---|---|---|---|
| **Branscheidt Exp1, 2019** | Sequential pinch-force task (40 % MVC) | maximal Isometric pinch until force decline below highest target force (40% MVC). Control group performed pinch at 5%MVC for the same time | Neuromuscular | Sequence Learning (Discrete), De Novo Learning (Continuous), Motor Acuity | 24h (Â±1h) |
| **Branscheidt Exp2, 2019** | Same pinch-force paradigm | maximal Isometric pinch until force decline below highest target force (40% MVC). Control group performed pinch at 5%MVC for the same time | Neuromuscular | Sequence Learning (Discrete), De Novo Learning (Continuous), Motor Acuity | _ |
| **Branscheidt Exp3, 2019** | Same pinch-force paradigm | maximal Isometric pinch until force decline below highest target force (40% MVC). Control group performed pinch at 5%MVC for the same time | Neuromuscular | Sequence Learning (Discrete), De Novo Learning (Continuous), Motor Acuity | 24h (±1h) |
| **Branscheidt Exp4, 2019** | 10-element sequence tapping | maximal Isometric pinch until force decline below highest target force (40% MVC). Control group performed | Neuromuscular | Sequence Learning (Discrete) | 24h (±1h) |

| | | | | | |
|---|---|---|---|---|---|
| | | pinch at 5%MVC for the same time | | | |
| **Carron, 1972** | Ladder-climb balance task | Cycle ergometer to ≥ 180 bpm (control: vowel detection) | Mixed (Cardiovascular, Neuromuscular) | Sequence Learning (Continuous), Motor Acuity | 24h |
| **Carron, 1969** | Pursuit rotor (stylus tracking) | Hand ergometer at 200 kgm/min until exhaustion | Mixed (Cardiovascular, Neuromuscular) | Motor Acuity | 24h |
| **Carron, 1971** | Stabilometer balancing (20 s trials) | Bike ergometer HR ≥ 180 bpm before & between trials; Session 3: no fatigue Control group cancelled vowels | Mixed (Cardiovascular, Neuromuscular) | Sequence Learning (Continuous), De Novo Learning (Continuous), Motor Acuity | 48h |
| **Cochran, 1975** | Stabilometer | Exp.: Bicycle ergometer pedaling at 600 kg·m/min, 50 rpm; Ctrl.: rest or puzzle between trials | | Sequence Learning (Continuous), De Novo Learning (Discrete), Motor Acuity | 1 week |
| **Cotten, 1974** | Volleyball mirror-toss (5 throws/trial for 50 throws) | 7 minutes of severe overall body exercise (steps; total body group, 30 ascent / minute)) vs 5 minutes of severe local exercise (biceps curl agains 23 pound, | Mixed (Cardiovascular, Neuromuscular) | Sequence Learning (Continuous), De Novo Learning (Discrete), Motor Acuity | 1 week |

| | | | | | |
|---|---|---|---|---|---|
| | | 30 rep per minute, pronated) + 30 sec of fatiguing exercise every 10 throws. Control group: rest | | | |
| **Cotten, 1972** | Mirror Target Toss (50 throws + 30 s exercise/rest) | Stool stepping 30 up/down per min for 7 min + 30 s every 2 trials | Mixed (Cardiovascular, Neuromuscular) | Sequence Learning (Continuous), De Novo Learning (Discrete), Motor Acuity | 24h |
| **Dickinson Exp1, 1979** | Fitts' reciprocal tapping | Arm cranking rotated the pedal of a Monark bicycle ergometer with either their preferred or non-preferred hand.at 40 rpm to exhaustion (4–7 min avg) | Mixed (Cardiovascular, Neuromuscular) | Motor Acuity | 24h |
| **Dwyer, 1984** | Ladder climb (Bachman) | Step-up activity: 60 times a minute onto a bench with a horizontal bar above the bench that subjects had to grab onto | Mixed (Cardiovascular, Neuromuscular) | Sequence Learning (Continuous), Motor Acuity | 24h |
| **Ferrer-Uris, 2018** | Rotational visuomotor adaptation (rVMA) | 13-min 20 m shuttle run alternating 3 min | Mixed (Cardiovascular, Neuromuscular) | Adaptation | 5 days |

| | | | | | |
|---|---|---|---|---|---|
| | | @85 % $VO_2max$/2 min @60 % $VO_2max$ | | | |
| **Goodwin, 1971** | Sigma task (arm ergometer) | Friction arm ergometer: 2 min @60 rpm + 15 s between trials | Cardiovascular | Sequence Learning (Discrete), Sequence Learning (Continuous), Motor Acuity | 1h, 24h, 7d |
| **Mierau, 2009** | Tracking with left–right visuomotor inversion | Incremental treadmill running (2.0 → +0.5 m/s every 5 min) until exhaustion | Mixed (Cardiovascular, Neuromuscular) | De Novo Learning (Continuous), Motor Acuity | 3 days |
| **Moghani, 2021** | Grip-force regulation (target vs produced force) | Stroop color-word test (1 250 responses in 60 min) | Cardiovascular | Motor Acuity | same session |
| **Nunney, 1963** | Snoddy stabilimeter & Rotary Pursuit | Bike ergometer/treadmill at various loads/gradients (4 groups) | Cognitive | Sequence Learning (Continuous), De Novo Learning (Continuous), Motor Acuity | 10 min, 20 min, 24h |
| **Pack, 1974** | Ladder climb (rungs per time) | Treadmill. Started at 3mph at 3% gradient. This was gradually increased until target heart rate was reached. When target HR was reached the | Mixed (Cardiovascular, Neuromuscular) | Sequence Learning (Continuous), Motor Acuity | 6 days acquisition, 48h spacing |

| | | | | | |
|---|---|---|---|---|---|
| | | intensity was maintained for 3 minutes so HR was maintained. | | | |
| **Schmidt, 1969** | Ladder climb (30 s per trial) | Bicycle ergometer: mild (750 kgm/min) vs moderate (1 200 kgm/min); control: vowel cancelling | Mixed (Cardiovascular, Neuromuscular) | Sequence Learning (Continuous), Motor Acuity | 24h |
| **Siekirk, 2019** | Proprioceptive matching (kinesthesiometer) | Elbow flexion 75–85 % max (6–10 reps) until exhaustion | Mixed (Cardiovascular, Neuromuscular) | Motor Acuity | 48h |
| **Stockard, 1974** | Multi-station circuit (balance, kick, catch) | Ergometer at 900 vs 600 kpm/min | Neuromuscular | Sequence Learning (Continuous), Motor Acuity | 48h |
| **Takahashi, 2006** | Force-field reaching (elastic bands pull) | Reaching against elastic bands until inability to meet 6 s time limit | Mixed (Cardiovascular, Neuromuscular) | Adaptation, Motor Acuity | 1 week |
| **Thomas, 1975** | Stabilometer balancing (frontal plane) | Treadmill walk to reach 180 bpm | | Sequence Learning (Continuous), De Novo Learning (Continuous), Motor Acuity | 10 min |
| **Wanner, 2020** | Stabilometer balancing (±20° platform) | High-intensity group (EX-H): ~90% Wmax in the “work” intervals. | Mixed (Cardiovascular, Neuromuscular) | Sequence Learning (Continuous), De Novo | 24h |

| | | | | | |
|---|---|---|---|---|---|
| | | Moderate-intensity group (EX-M): ~45% Wmax in the “work” intervals. Low-intensity (EX-L): 25 W continuous (minimal). | | Learning (Continuous), Motor Acuity | |
| **Williams, 1976a** | Ladder climb (Bachman) | Step-up 30/min (22″ bench) + arm use, maintain HR ≥ 180 bpm | Mixed (Cardiovascular, Neuromuscular) | Sequence Learning (Continuous), Motor Acuity | 24h |
| **Williams, 1976b** | Two-hand pursuit rotor | 4 min step-up to HR + 1 min step-up between each 1 min tracking block; Day 2: no fatigue | Mixed (Cardiovascular, Neuromuscular) | Motor Acuity | 24h |
| **Williams, 1975** | Hand-crank pursuit rotor | Hand-cranking ergometer; control: backward count | Mixed (Cardiovascular, Neuromuscular) | Motor Acuity | 48h |
| **Zabihhosseinian, 2020** | Sinusoidal tracing (touchpad) | Cervical extensor hold with 2 kg until > 5 s deviation | Neuromuscular | Motor Acuity | 24h |
| **Zabihhosseinian, 2021** | Sinusoidal tracing with color feedback | Cervical extensor hold with 2 kg until > 5 s deviation | Neuromuscular | Motor Acuity | 24h |
